# Similarities and differences between native HIV-1 envelope glycoprotein trimers and stabilized soluble trimer mimetics

**DOI:** 10.1101/500975

**Authors:** Alba Torrents de la Peña, Kimmo Rantalainen, Christopher A. Cottrell, Joel D. Allen, Marit J. van Gils, Jonathan L. Torres, Max Crispin, Rogier W. Sanders, Andrew B. Ward

## Abstract

The HIV-1 envelope glycoprotein (Env) trimer is located on the surface of the virus and is the target of broadly neutralizing antibodies (bNAbs). Recombinant native-like soluble Env trimer mimetics, such as SOSIP trimers, have taken a central role in HIV-1 vaccine research aimed at inducing bNAbs. We therefore performed a direct and thorough comparison of a full-length native Env trimer containing the transmembrane domain and the cytoplasmic tail, with the sequence matched soluble SOSIP trimer, both based on an early Env sequence (AMC011) from an HIV^+^ individual that developed bNAbs. The structures of the full-length AMC011 trimer bound to either bNAb PGT145 or PGT151 were very similar to the structures of SOSIP trimers. Antigenically, the full-length and SOSIP trimers were comparable, but in contrast to the full-length trimer, the SOSIP trimer did not bind at all to non-neutralizing antibodies, most likely as a consequence of the intrinsic stabilization of the SOSIP trimer. Furthermore, the glycan composition of full-length and SOSIP trimers was similar overall, but the SOSIP trimer possessed slightly less complex and less extensively processed glycans, which may relate to the intrinsic stabilization as well as the absence of the membrane tether. These data provide insights into how to best use and improve membrane-associated full-length and soluble SOSIP HIV-1 Env trimers as immunogens.

## Introduction

The HIV-1 envelope glycoprotein (Env) trimer is the target of broadly neutralizing antibodies (bNAbs) that arise during HIV-1 infection and is, therefore, an attractive immunogen for vaccine design. Previous studies have reported that bNAbs provide passive protection from viral challenges in macaques [1–3]. One approach to induce protective bNAbs is the use of Env-based vaccines that mimic native Env on the virus.

Previously, we described a soluble Env trimer, BG505 SOSIP.664, which contains an I559P substitution in gp41 that stabilizes the prefusion state of Env and a disulfide bond that covalently links the two subunits of the Env protein, gp120 and gp41 [4–6]. Subsequently, soluble SOSIP trimers from different clades have been described and characterized biophysically and biochemically [4,7–11]. The high resolution structures of several of these SOSIP trimers were solved by cryo-electron microscopy (cryo-EM) and x-ray crystallography, allowing for structure-based Env trimer vaccine design [7,11–17]. Soluble SOSIP Env trimers have been tested as immunogens in animals and elicited neutralizing antibody (NAb) responses against the autologous viruses but generally not against heterologous Tier-2 viruses [10,18–22].

Native-like soluble Env trimers, such as SOSIP trimers, lack the membrane proximal external region (MPER), a target for broadly neutralizing antibodies, the transmembrane domain (TM), and the cytoplasmic tail. It is not entirely clear whether or how the absence of these domains and the presence of the SOSIP mutations influence the detailed properties of the trimers. Cryo-EM studies showed that a membrane-derived trimer that lacked the cytoplasmic tail and that was bound to bNAb PGT151 closely resembled SOSIP trimers at the structural level [23], but the lack of sufficient amounts of protein prevented a detailed antigenic and glycan characterization of these membrane-derived trimers. There is only limited data available comparing the antigenic and biophysical properties of SOSIP trimers versus corresponding full-length trimers because full-length trimers, as well as virion-derived trimers, are difficult to purify in sufficient quantities to perform such comparative studies.

In this study we compared the structural, biochemical and biophysical properties of a highly expressed full-length Env trimer from an elite neutralizer who was infected with a subtype B virus (AMC011) [24,25], with those of the corresponding sequence matched AMC011 SOSIP trimer. Using cryo-EM we generated structural models of the full-length AMC011 Env trimer bound to either the PGT145 Fab or the PGT151 Fab. Both of these quaternary specific bNAbs bind to and stabilize a similar conformation that resembles all previously reported membrane-derived and soluble SOSIP Env structures. Furthermore, we investigated the composition of the glycan shield of full-length and SOSIP trimers by site-specific glycosylation profiling using mass spectrometry, and the antigenic structure by bio-layer interferometry (BLI; Octet), flow cytometry and neutralization assays. The results show that the antigenic profile and glycan composition of full-length and SOSIP trimers was similar, but also revealed subtle and interesting differences. In a similar manner to previous observations comparing virally derived N-glycans and corresponding SOSIP trimers [26,27], there was a noticeable decrease in the number and processing of complex-type glycans on the SOSIP trimer. This most likely can be attributed to its enhanced stability and reduced conformational breathing due to the presence of the SOSIP mutations, as well as the lack of the membrane tether. Furthermore, in contrast to the full-length AMC011 trimer, the corresponding SOSIP trimer did not bind to a number of non-neutralizing antibodies (non-NAbs), which probably also relates to the enhanced stability and reduced sampling of alternative conformations. These results should guide the use and improvement of full-length and soluble Env trimers as vaccine immunogens.

## Results

### Purification of full-length Env trimers

To allow for a direct structural, antigenic and biophysical comparison between SOSIP.664 and full-length trimers, we selected the AMC011 clade B *env* gene, which is a consensus sequence of early Env sequences from an HIV-infected individual enrolled in the Amsterdam Cohort Studies on HIV/AIDS (ACS) [24,25].

The selection of the AMC011 sequence was based on three criteria. First, the patient that was infected with the AMC011 virus developed bNAbs and qualified as an elite neutralizer [24,25]. As such, this particular Env is of relevance for vaccine design aimed at inducing bNAbs. The ACS202 bNAb that was isolated from this patient targets the fusion peptide and the gp120-gp41 interface [24]. Second, the neutralization sensitivity of the AMC011 virus places it in the grey area between the Tier-1B and Tier-2 categories of neutralization sensitivity. While the AMC011 virus was initially categorized as a Tier-2 virus, typical of circulating difficult to neutralize viruses [24,28], follow-up experiments using a broader set of antibodies and plasma reagents revealed that the AMC011 virus could be neutralized by some V3 and CD4-targeting antibodies, placing it in the Tier-1B category (S1 Table). Because neutralization sensitivity relates to conformational breathing of the Env trimer [28,29], we argued that the Tier-1B status of the AMC011 virus would allow us to observe a discernable impact of SOSIP stabilization. Third, the full-length AMC011 trimer was expressed at high yields (see below), making a thorough biophysical characterization feasible from a practical perspective.

Using a previously described procedure for the purification of native Env trimers from membranes [23], we isolated native full-length AMC011 Env bound to PGT151 Fab or PGT145 Fab from the surface of HEK293F cells that were transiently transfected with expression vectors for the full-length AMC011 gp160 and furin. bNAbs PGT145 and PGT151 both require a native quaternary structure, recognizing the trimer apex and fusion peptide-gp120-gp41 interface, respectively. In contrast to previous experiments using different Env sequences that had resulted in only modest yields of purified full-length Env (between ∼45 and ∼75 μg/L) [30], we successfully produced full-length AMC011 trimers at yields of ∼350 μg/L using PGT151 Fab and ∼65 μg/L using PGT145 Fab (Figs 1A, 1B and S1A). For comparative experiments, we also purified soluble AMC011 SOSIP.664 trimers with PGT151 or PGT145 affinity chromatography columns, as previously described (Fig 1B) [10,31].

**Figure 1.**
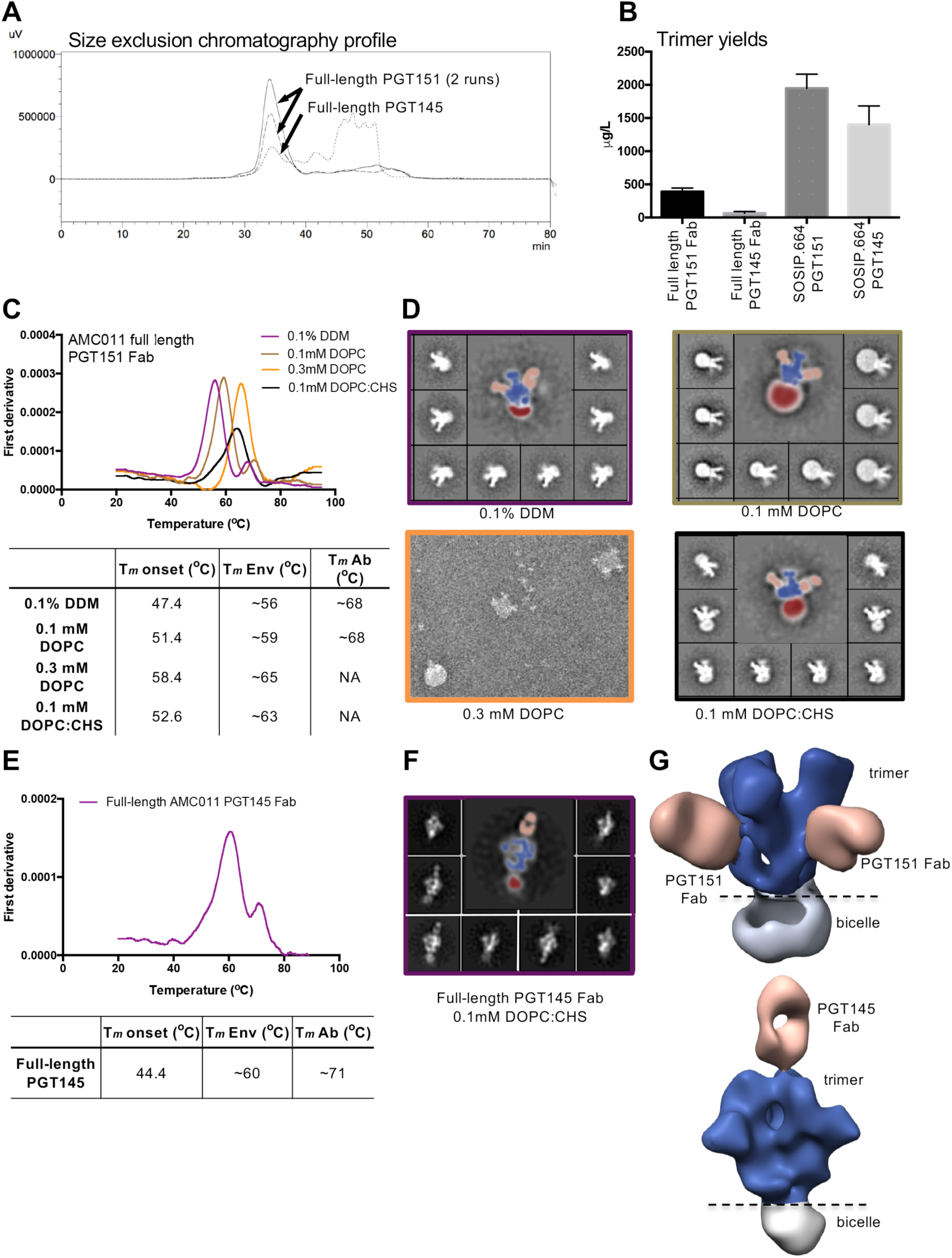
Purification of the full-length AMC011 trimer in complex with PGT151 or PGT145 Fab. (**A**) Size exclusion chromatography profiles of full-length AMC011 Env bound to PGT151 Fab (two runs) or bound to PGT145 Fab (one run). (**B**) Average yields of full-length Env constructs bound to PGT151 Fab or PGT145 Fab and their SOSIP.664 counterparts. For comparison, soluble AMC011 SOSIP.664 trimers were purified with PGT151 and PGT145 affinity chromatography columns. The mean yields shown in the graph represent the averages of three to six purifications. (**C**) Thermostability of full-length AMC011 trimers in DDM or at different concentrations of DOPC and DOPC:CHS (1:1 molar ratio) after detergent removal. The unfolding pattern of the different proteins was assessed by plotting the first derivative of the curves acquired with nano-DSF. The *T*_m_ onset, the melting temperatures of the Env (*T*_m_ Env) and the *T*_m_ of the antibody (*T*_m_ Ab) are shown in the table. The averages *T*_m_ values derived from two to five different experiments are listed in the table. The raw data of one such experiment is depicted in the graph. NA: not applicable. (**D**) 2D class averages from negative-stain EM images are shown for the full-length AMC011 Env trimer bound to PGT151 Fab in DDM or at different concentrations of DOPC or DOPC:CHS (1:1 molar ratio). Colors in the thermostability graph match the 2D class average boxes. (**E**) Thermostability of full-length AMC011 Env trimer bound to PGT145 Fab in 0.1 mM DOPC:CHS (1:1 molar ratio). The *T*_m_ was calculated from one experiment. (**F**) Representative 2D class averages from full-length AMC011 Env trimer in 0.1 mM DOPC:CHS. (**G**) 3D reconstructions of full-length AM011 trimer bound to PGT151 or to PGT145, which were derived from the class-averages shown in (D) and (F). Coloring in (D), (F) and (G) is as follows: PGT151 and PGT145 in light orange, the bicelle in gray and the envelope trimer in dark blue.

Next, we modified the cryo-EM protocol that was optimized for the JR-FL Env trimer lacking the cytoplasmic (ΔCT) [23] to be applicable for the full-length AMC011 Env trimer, which was reconstituted in a homogeneously shaped bicelle. Specifically, full-length AMC011 trimers bound to the PGT151 Fab were solubilized from the cell membrane TX-100 detergent micelles, followed by exchange into n-Dodecyl-β-D-maltoside (DDM) micelles and finally reconstituted reconstituted into a lipid bicelle containing different concentrations of 1,2-dioleoyl-*sn*-glycero-3-phosphocoline (DOPC). While the incorporation of 0.3 mM of DOPC led to the formation of liposomes containing full-length Env, the incorporation of 0.1 mM of DOPC facilitated the formation of bicelles with different sizes (Figs 1C and 1D). Next, we used DOPC in a 1:1 molar combination with cholesteryl hemisuccinate (CHS), which resembles the lipid composition of microdomains present at the cell membrane of infected CD4+ T cells [32]. The detergent exchange from DDM to DOPC:CHS of the full-length Env trimer sample led to the generation of homogeneously shaped bicelles, which surround the transmembrane domains of the Env protein and increased the stability of the full-length Env trimer, as shown by an elevation of the midpoint of thermal denaturation (*T*_m_) from ∼56.1°C to ∼63.1°C, reaching similar stability levels as soluble AMC011 SOSIP.664 (*T*_m_ value of 61.8°C) (Figs 1C, 1D and S1B). Reconstitution of the full-length Env trimer bound to PGT145 in DOPC:CHS led to the formation of similarly homogeneous shaped bicelles and also increased thermostability (*T*_m_ value of ∼60.4°C) (Figs 1E and 1F). Finally, negative stain (NS)-EM reconstructions of the trimer Fab complexes in DOPC:CHS showed that both bNAbs PGT151 and PGT145 bound to the full-length Env trimer in an asymmetric manner as previously described, with two and one Fab per trimer, respectively (Fig 1G) [33,34]. Overall, while the wild type, full-length Env trimers are inherently unstable and cannot be purified easily, the addition of the quaternary PGT151 or PGT145 Fabs together with the use of DOPC and CHS, resulted in the purification of stable complexes.

The AMC011 SOSIP.664 trimer, which was purified with the PGT151 affinity chromatography column, displayed *T*_m_ value of 61.8°C. The trimer was however more stable than the full-length trimer when analyzed in complex with PGT151 or PGT145 (*T*_m_ values of 73.6°C and 67.0°C, respectively; increases of 10.5°C and 6.6°C compared to the full-length trimer in complex with PGT151 or PGT145; increases of 11.8°C and 5.2°C compared to SOSIP.664 alone) (S1B Fig). We compared the conformation of the full-length and soluble SOSIP.664 trimers by negative-stain (NS)-EM and found that all complexes yielded similar 2D class-averages, suggesting that neither antibody nor the presence or absence of the bicelle biased the trimer conformation (Figs 1D, 1F and S1B).

### Antigenic comparison of AMC011 SOSIP.664 and full-length trimers

BLI was used to assess binding of a panel of well-characterized bNAbs and non-NAbs (n=21 in total) to the purified SOSIP.664 and full-length trimers in solution. We defined non-NAbs as antibodies that cannot neutralize typical neutralization-resistant Tier-2 viruses. Such non-NAbs might, however, neutralize Tier-1 viruses, and in fact the Tier-1B virus AMC011 is neutralized by some of these non-NAbs (see below and Table S1). We also note that the BLI experiments were carried out with unliganded SOSIP.664 trimers (purified with PGT151 affinity chromatography column) and full-length trimers bound to either PGT145 Fab or PGT151 Fab, resulting in competition effects when analyzing the binding of mAbs to overlapping epitopes [35]. Overall, we observed that full-length and SOSIP.664 trimers were very similar antigenically, with a few interesting exceptions that are specified below.

First, we determined the binding of quaternary antibodies that target the apex of the trimer (PGT145, PGDM1400 and PG16)[34,36]. All three bNAbs bound to the full-length trimer that was bound to PGT151 Fab, but as expected PGT145 did not bind to the trimer that was already bound to PGT145 (Figs 2A and S2). While PGT145 and PGDM1400 showed increased recognition of the SOSIP.664 compared to the full-length trimer, PG16 showed higher affinity for the full-length trimer (Fig 2A).

**Figure 2.**
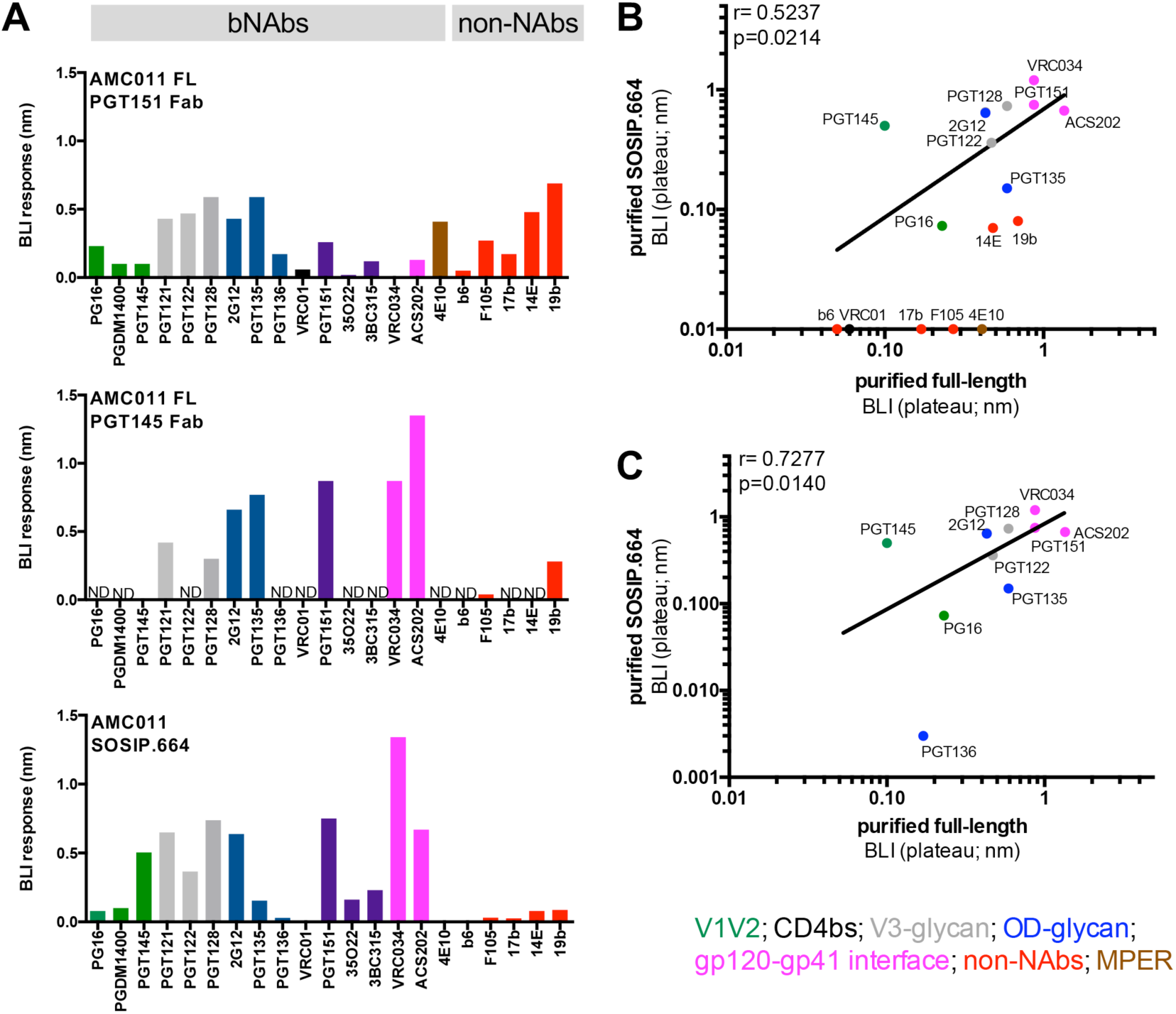
Antigenicity profile of purified full-length AMC011 Env trimer determined by BLI. (**A**) The Antigenicity of full-length and SOSIP.664 Env trimers was assessed by BLI (Octet). Bars represent the maximum values of the binding curves (Fig S2). A panel of 16 bNAbs and 5 non-NAbs was tested: apex directed antibodies (green), V3-glycan (blue), CD4 binding site (black), gp120-gp41 interface (magenta), MPER (brown) and non-NAbs (red). ND: not determined (the lower yields of full-length AMC011 Env in complex with PGT145 precluded a full analysis). (**B**) The binding of bNAbs and non-NAbs to full-length and to SOSIP.664 Env trimers, assessed by BLI, was compared. The Spearman correlation (inset) is shown. (**C**) Same analysis as in (B), but excluding the non-NAbs and MPER-directed NAb. Antibodies are colored as in panel (A).

We also determined binding of bNAbs that target the outer-domain and the V3 glycan region of the Env trimer [37–40]. While PGT121, PGT122 and PGT128, which target the N332 glycan outer domain region, bound similarly to the full-length and the SOSIP.664 trimer, PGT135 and PGT136, which also target the N332 glycan, but from a different angle of approach, showed an increased binding to full-length compared to SOSIP.664 trimers (Fig 2A). The presence of different glycoforms at positions N386 and N392 might explain these differences, as discussed below [38,41].

Next, we tested the binding of the broadly neutralizing antibody isolated from the AMC011 elite neutralizer, ACS202, which targets the fusion peptide and the glycan at position 88 [24]. As expected, ACS202 and other antibodies that target the interface of the Env trimer (PGT151, 35O22 and VRC34) did not bind efficiently to the full-length trimer that was already bound to PGT151 Fab. However, ACS202, PGT151 and VRC34 interacted strongly with the PGT145 bound full-length trimer. Furthermore, ACS202 bound to the full-length Env trimer with a higher on-rate than to the SOSIP.664 (Figs 2A and S2). Since this antibody binds to the glycan at position N88, this difference could be explained by the presence of different glycoforms at position N88 on full-length and SOSIP.664 trimers (see below). On the other hand, VRC34, which also targets the fusion peptide and the N88 glycan [42], showed similar binding to both SOSIP.664 and full-length trimer (Fig 2A).

The membrane proximal external region (MPER) is only present in the full-length Env trimer. Hence, the MPER-directed antibody 4E10 only showed binding to the full-length Env trimer and not to the SOSIP.664 trimer (Fig 2A).

Finally, we also tested the binding of five antibodies directed to the CD4 binding site (CD4bs; b6 and F105), the CD4-induced site (CD4i; 17b) and the V3 (14e and 19b) that typically only neutralize Tier-1 viruses (easy to neutralize viruses) and that we term non-NAbs. Except for b6, all these non-NAbs bound to the full-length AMC011 trimer by BLI, a finding that is probably related to the Tier-1B status and inherent conformational flexibility. Conversely, the non-NAbs did not bind to the SOSIP.664 trimer.

We compared the antigenicity of full-length trimers bound to either PGT145 Fab or PGT151 Fab. For comparison, we omitted the antibodies that competed with the Fabs bound to the full-length trimers because of overlapping epitopes (i.e. PG16, PGDM1400 and PGT145 for Env bound to PGT145 Fab, and VRC34, ACS202, 3BC315 and 35O22 for Env bound to PGT151 Fab). A correlation between the BLI values against the two full-length trimers was observed (Spearman correlation coefficient r=0.7414, p=0.0223, S3 Fig), indicating that full-length trimers purified in complex with PGT145 or PGT151 Fab showed similar antigenic profiles. When full-length and SOSIP.664 AMC011 trimers were compared, the correlation was weak (r=0.5237, p=0.0214; Fig 2B), likely the result of binding of the non-NAbs and the MPER-directed 4E10 bNAb to full-length, but not to SOSIP.664 Env trimers. Indeed, when the non-NAbs and 4E10 were excluded, the correlation between binding to full-length and SOSIP.664 trimers was strong (r=0.7277, p=0.0140; Fig 2C).

To provide a frame of reference for these BLI data, we assayed the same panel of bNAbs and non-NAbs in neutralization assays with the parental AMC011 virus, and performed binding assays to full-length AMC011 trimers on the surface of transfected cells by FACS analysis (Fig 3A). To correlate parameters from the different assays we used maximal response values derived from BLI assays (plateau) and area under the curve (AUC) values from FACS assays and neutralization assays. A very strong correlation was observed when comparing antibody binding to membrane-bound AMC011 trimers by FACS analysis *versus* AMC011 virus neutralization in TZM-bl assays (r=0.8343, p<0.0001, Fig 3B), indicating that prior to purification from the membrane, the full-length AMC011 Env trimer was a close mimic of the functional AMC011 trimer on the virus.

**Figure 3.**
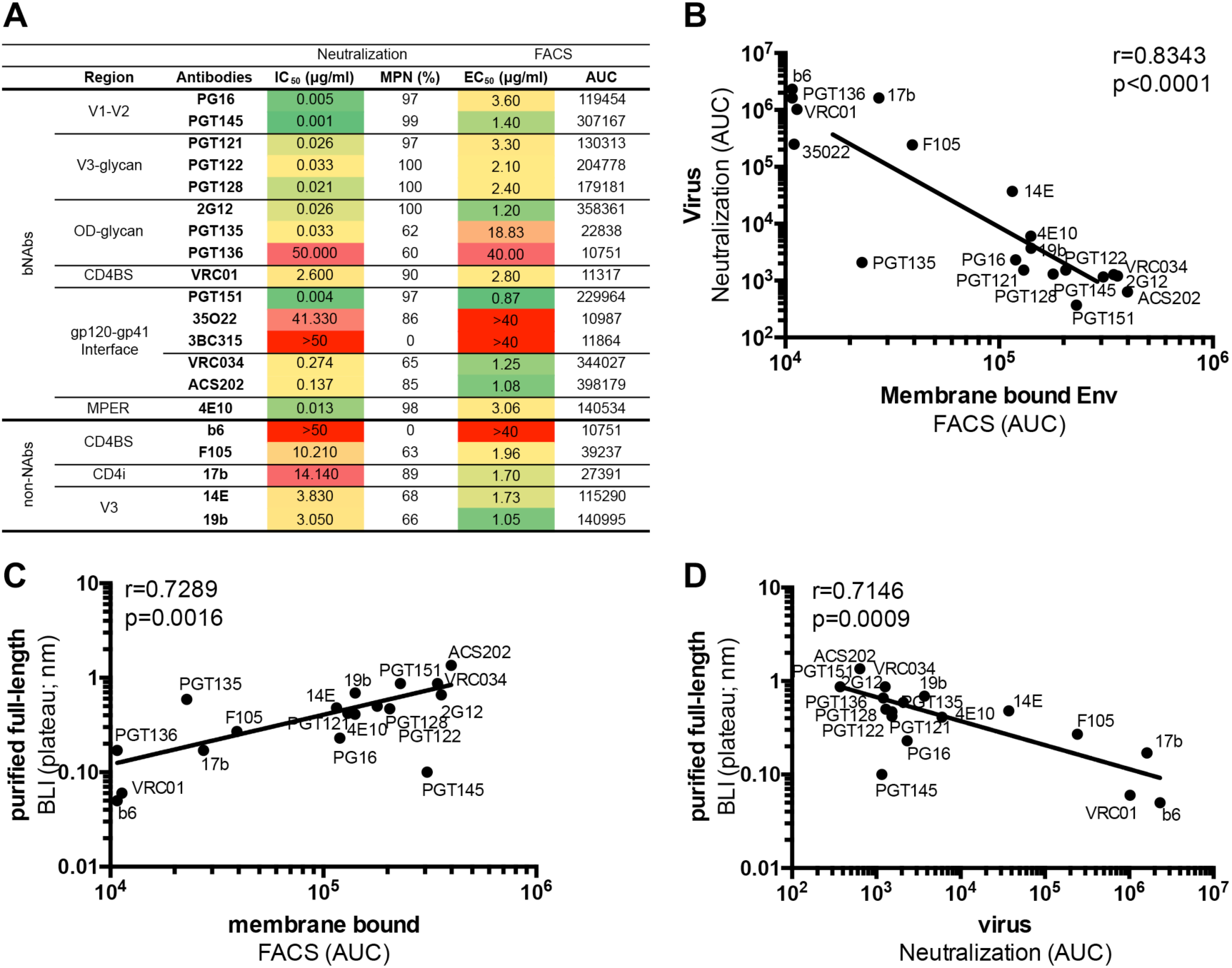
Antigenicity profile of the full-length Env protein present on the cell membrane or on the virus. (**A**) Neutralization of the AMC011 virus was assessed as a proxy for antibody binding to the functional AMC011 trimer on the virus. IC_50_ and maximum percentage neutralization (MPN) values for a panel of bNAbs and non-NAbs against AMC011 virus are shown. Binding of bNAbs and non-NAbs to the Env protein embedded in the membrane of HEK293F cells was determined by FACS. Half maximal binding concentrations (EC_50_) and area under the curve (AUC) values are shown. Data are representative for two to four different experiments. (**B**) Comparison of AMC011 antigenicity obtained by FACS (membrane-bound Env) and neutralization assays (virus-associated functional Env), by FACS and BLI (purified membrane-derived Env) (**C**) and by neutralization assays and BLI (**D**) are shown.

We next investigated how antibody binding to the purified full-length AMC011 trimer, as assessed by BLI, compared to binding to the cell-associated trimer and the virus-associated trimer as assessed by FACS (EC_50_) and neutralization (IC_50_), respectively. Strong correlations were observed between antibody reactivity assessed by BLI and FACS, and BLI and neutralization (r=0.7289, p=0.0016 and r=0.7146, p=0.0009; Figs 3C and 3D). We therefore conclude that the full-length membrane-derived and purified AMC011 trimers, the unpurified trimers on the surface of HEK293F cells and the functional trimers on the AMC011 virus, adopt very similar antigenic structures. In line with the BLI data, four of five non-NAbs tested (F105, 17b, 19b, 14e, but not b6) neutralized the virus and bound the full-length Env trimer in neutralization assays and FACS, respectively (Figs 3A, S4 and S5). This observation is likely due to the conformational plasticity of the native, non-SOSIP stabilized full-length Env trimer, in line with the Tier-1B status. The AMC011 virus, being a Tier-1B virus probably samples conformations that transiently expose the CD4bs, CD4i and V3 epitopes of these non-NAbs.

The corresponding AMC011 SOSIP.664 trimers, on the other hand, did not show appreciable binding to the non-NAbs (Fig 2A). Hence, the SOSIP mutations resulted in desirable stabilization of the prefusion Env trimer that had a decreased propensity to sample alternative conformations and expose undesirable non-NAb epitopes.

### Structural comparison of full-length AMC011 trimers bound to PGT145 or PGT151 and SOSIP trimers

We generated structures of the full-length AMC011 trimer bound to PGT151 or PGT145 Fab by single particle cryo-EM at resolutions of ∼4.2 Å and ∼5.7 Å, respectively (Figs 4A, Table 1 and S6A). We note that we could not reconstruct the MPER, TM and CT domains on the full-length AMC011 trimers, probably because of the flexible nature of these domains in the bicelle milieu. Therefore, the following comparison was performed using the ectodomains only (up to residue 664).

**Figure 4.**
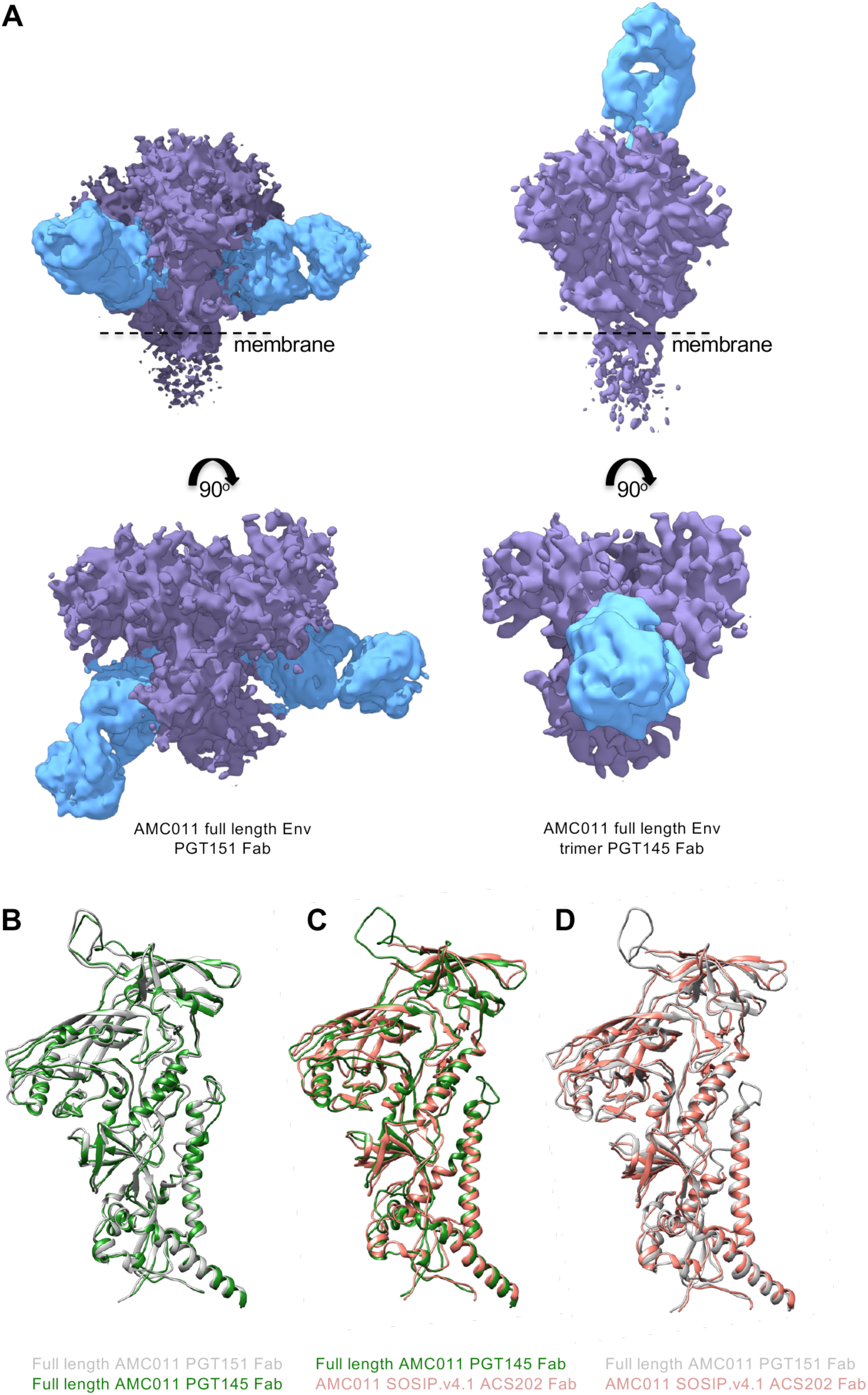
Cryo-EM structure of full-length AMC011 Env in complex with PGT145 Fab or PGT151 Fab and comparison with other structures. **(A)** The cryo-EM map of the full-length AMC011 trimer bound to PGT151 Fab (left), the reconstruction of which is at 5.0 Å and the cryo-EM map of the full-length AMC011 trimer bound to PGT145 Fab (right), the reconstruction of which is at 5.7 Å are shown. Superimposition of the full-length AMC011 Env bound to PGT145 Fab (in green) with: (**B**) full-length AMC011 bound to PGT151 Fab (in gray) and (**C**) AMC011 SOSIP.v4.2 (in light red). Superimposition of the full-length AMC011 Env bound to PGT151 Fab (in gray) with: (**D**) AMC011 SOSIP.v4.2 (in light red).

**Table 1.**
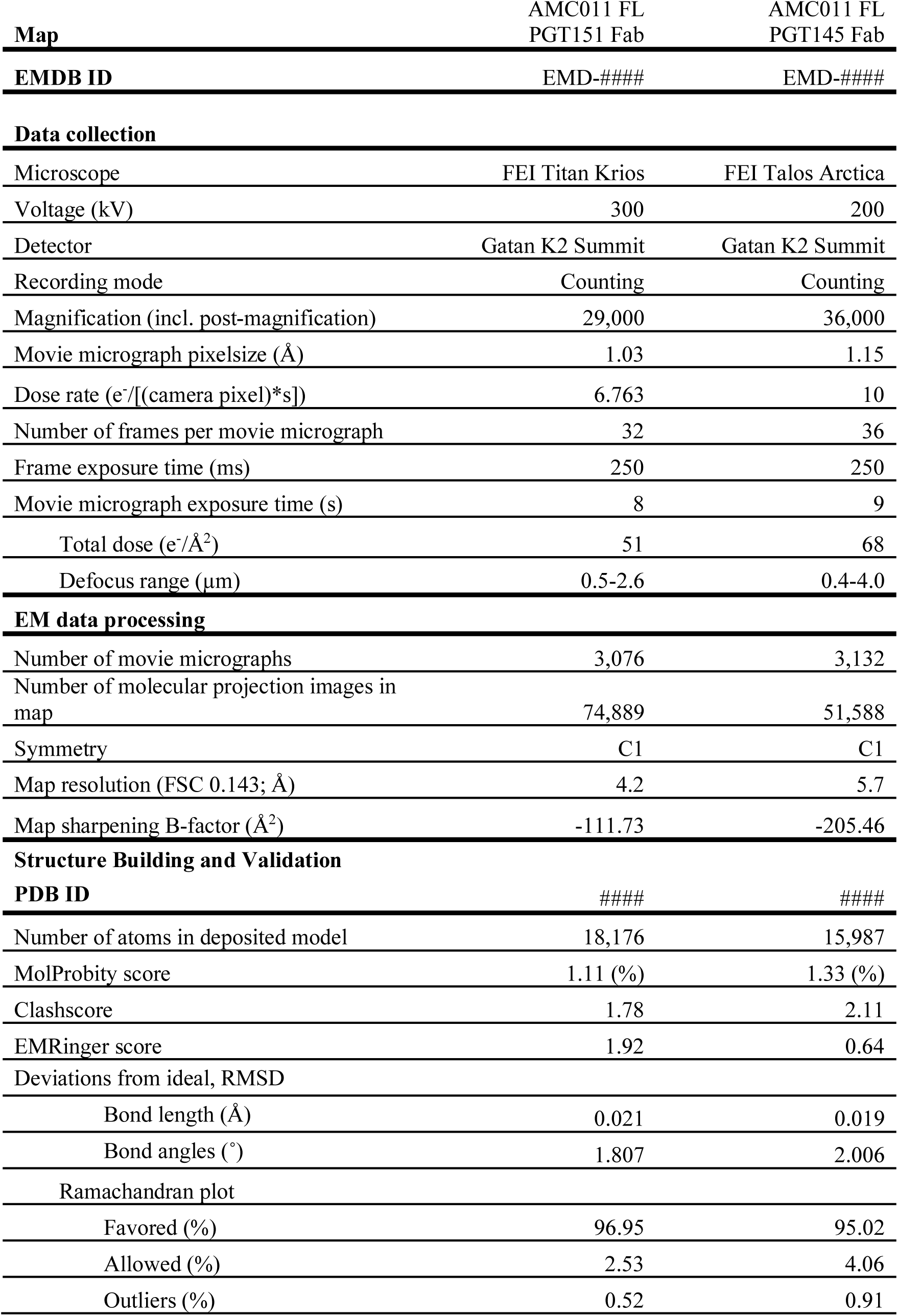
EM data collection and refinement statistics

| Map | AMC011 FL<br>PGT151 Fab | AMC011 FL<br>PGT145 Fab |
| --- | --- | --- |
| EMDB ID | EMD-#### | EMD-#### |
| **Data collection** |  |  |
| Microscope | FEI Titan Krios | FEI Talos Arctica |
| Voltage (kV) | 300 | 200 |
| Detector | Gatan K2 Summit | Gatan K2 Summit |
| Recording mode | Counting | Counting |
| Magnification (incl. post-magnification) | 29,000 | 36,000 |
| Movie micrograph pixelsize (Å) | 1.03 | 1.15 |
| Dose rate (e<sup>-</sup>/[(camera pixel)*s]) | 6.763 | 10 |
| Number of frames per movie micrograph | 32 | 36 |
| Frame exposure time (ms) | 250 | 250 |
| Movie micrograph exposure time (s) | 8 | 9 |
| Total dose (e<sup>-</sup>/Å<sup>2</sup>) | 51 | 68 |
| Defocus range (μm) | 0.5-2.6 | 0.4-4.0 |
| **EM data processing** |  |  |
| Number of movie micrographs | 3,076 | 3,132 |
| Number of molecular projection images in map | 74,889 | 51,588 |
| Symmetry | C1 | C1 |
| Map resolution (FSC 0.143; Å) | 4.2 | 5.7 |
| Map sharpening B-factor (Å<sup>2</sup>) | -111.73 | -205.46 |
| **Structure Building and Validation** |  |  |
| PDB ID | #### | #### |
| Number of atoms in deposited model | 18,176 | 15,987 |
| MolProbity score | 1.11 (%) | 1.33 (%) |
| Clashscore | 1.78 | 2.11 |
| EMRinger score | 1.92 | 0.64 |
| Deviations from ideal, RMSD |  |  |
| Bond length (Å) | 0.021 | 0.019 |
| Bond angles (°) | 1.807 | 2.006 |
| Ramachandran plot |  |  |
| Favored (%) | 96.95 | 95.02 |
| Allowed (%) | 2.53 | 4.06 |
| Outliers (%) | 0.52 | 0.91 |

First, we assessed the similarities between the AMC011 trimer bound to the PGT151 Fab and to the PGT145 Fab. The two full-length AMC011 trimer structures were very similar to one another (backbone Cα r.m.s.d. between the gp120 domains is ∼1 Å and between the gp41 domains is ∼1 Å; Fig 4A), confirming that the structure was not altered to any large extent by the presence of either quaternary bNAb. The two structures also shared a highly similar architecture with the JR-FL ΔCT trimer (backbone Cα r.m.s.d. between the gp120 domains is ∼1 Å and between the gp41 domains is <1 Å for the comparison of AMC011 trimer bound to PGT151 with JRFL ΔCT trimer bound to PGT151; backbone Cα r.m.s.d. between the gp120 domains is ∼1 Å and between the gp41 domains is ∼1 Å for the comparison of AMC011 trimer bound to PGT145 with JRFL ΔCT trimer bound to PGT151; S6C and S6D Figs)[23]. Despite the low resolution of the previously solved structure of the AMC011 SOSIP.v4.2 trimer (6.2 Å)[24], both full-length structures appeared similar in conformation (backbone Cα r.m.s.d. between the gp120 domains is ∼1.2 Å and between the gp41 domains is ∼1 Å; Figs 4B and 4C).

The presence of PGT151 or PGT145 in the structures allowed us to investigate the binding interaction between the AMC011 trimer and the two bNAbs in detail. We compared the structure of the full-length AMC011 Env trimer in complex with PGT151 with that of JR-FL Env in complex with the same antibody, Both structures were similar to previous Env structures [14,23] (Fig 5A). The glycans belonging to the PGT151 epitope were ordered, and branched complex glycans at positions N611 and N637 in gp41 were resolved (Fig 5B), as observed in Lee et al., 2016.

**Figure 5.**
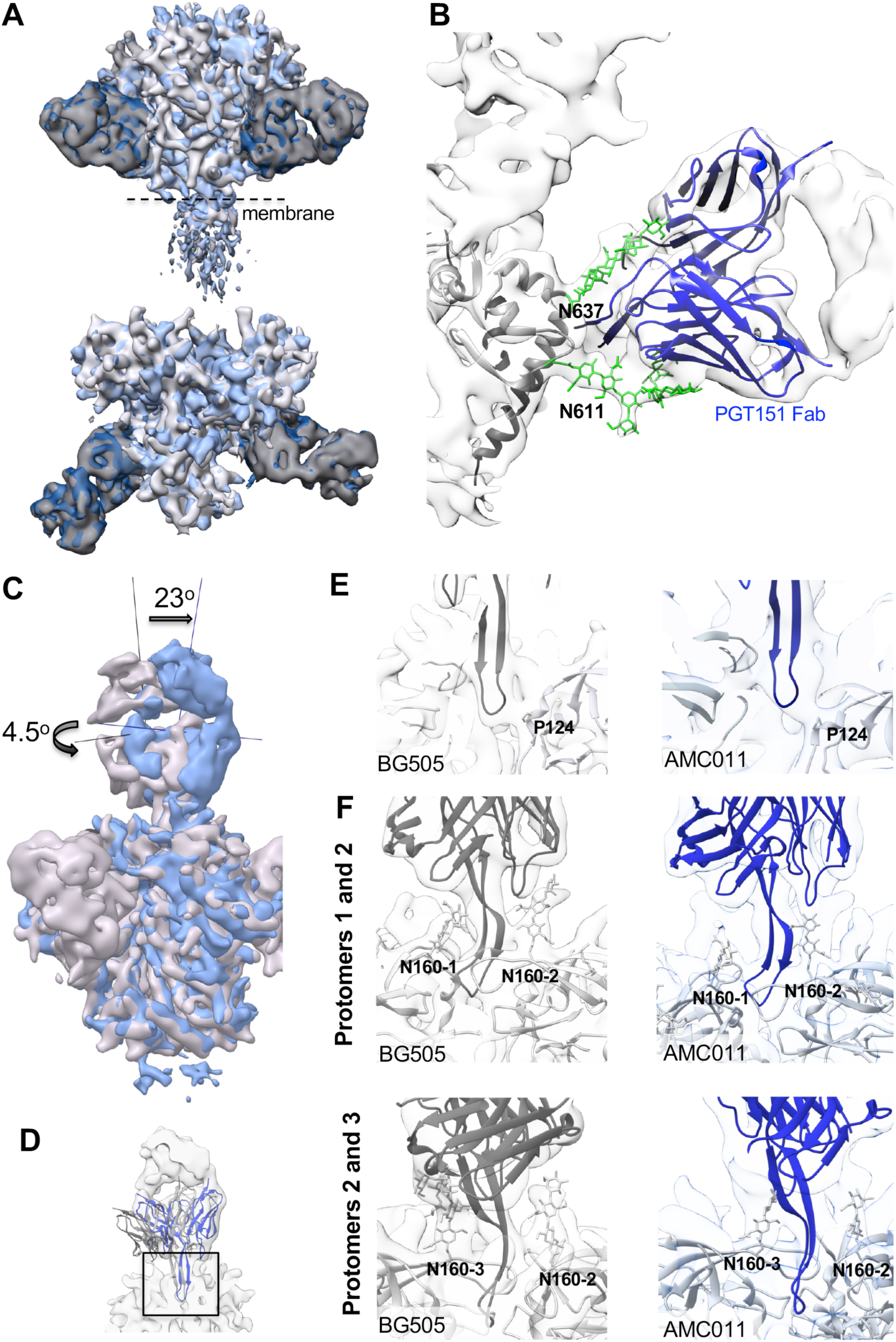
Binding of PGT145 Fab or PGT151 Fab to full-length AMC011 trimer. (**A**) Superimposition of the cryo-EM map of the full-length AMC011 Env bound to PGT151 Fab (blue) and the JR-FL ΔCT trimer bound to PGT151 Fab low-pass filtered at 5.0 Å (gray, Lee et al., 2016). (**B**) Close-up of the PGT151 epitope. Coloring is as follows: glycans are depicted in green, PGT151 Fab in blue and the cryo-EM reconstruction density in white. (**C**) Superimposition of the cryo-EM map of AMC011 full-length trimer bound to PGT145 and BG505 SOSIP.664 bound to PGT145 low-pass filtered to 5.7 Å. The vertical and horizontal axes were used to compare tilt and rotation angle between both PGT145 Fabs. (**D**) Close-up of the PGT145 epitope. The crystal structure of the PGT145 Fab (PDB: 3U1S) was docked into both cryo-EM maps for epitope comparison. The cryo-EM map of full-length AMC011 trimer bound to PGT145 is shown in white. (**E**) Cross sections of the PGT145 epitope (box in (D)) in BG505 SOSIP (gray) and AMC011 full-length (blue) trimers. Residue Pro124 of the trimer is shown in stick as a direct contact with the Fab. (**F**) Distribution of N160 glycans in the PT145 epitope of BG505 SOSIP and AMC011 full-length trimer. PGT145 Fab crystal structure (dark gray and dark blue for BG505 and AMC011, respectively) was docked into cryo-EM maps of BG505 SOSIP (EMD-8643) and AMC011 full-length trimers. The first sugars of the N160 glycan are indicated in sticks.

We next compared the structure and orientation of PGT145 bound to full-length AMC011 trimer and PGT145 bound to the BG505 SOSIP.664 trimer [34]. Interestingly, when the Env trimers were aligned we found that the PGT145 Fab was tilted by 23° and rotated by 4.5° when bound to the AMC011 trimer compared to when bound to the BG505 SOSIP.664 trimer (Fig 5C) [34]. Therefore, the trimer and PGT145 Fab were fit independently into the AMC011-PGT145 map to generate a pseudo-atomic model (Figs 5D and S6E). Despite this angular difference the location of the HCDR3 at the trimer 3-fold axis and the overall epitope was similar in both complexes (Figs 5E and 5F). The comparison suggests that either apex bNAbs approach different Env trimers at subtly different angles, or the SOSIP stabilizing mutations induce a more upright angle of approach.

Interestingly, during our 3D classification of the latter cryo-EM dataset we observed a class, representing ∼2 percent of particles, of dimeric full-length AMC011 Env bound to PGT145 and obtained a ∼10.2 Å resolution reconstruction of this class (middle panel in S7A Fig). This AMC011 dimer structure fits well into the structure of the AMC011 trimer bound to PGT145 (map to map correlation=0.9635), which is consistent with the previous finding that PGT145 strongly interacts with two protomers (S7B Fig) [34]. This class of AMC011 dimers was excluded from the trimer reconstructions.

### Composition of individual glycans on full-length and SOSIP trimers

Because glycans play important roles in the antigenicity and immunogenicity of Env, we next investigated the overall *N*-linked glycan profile of the soluble SOSIP.664 trimer and the full-length Env trimer bound to PGT151 Fab by using hydrophilic interaction liquid chromatography-ultra-performance liquid chromatography (HILIC-UPLC) [43]. While the full-length trimer contained 47% complex glycans, only 33% of the glycans were of the complex-type on SOSIP.664 (Fig 6A). This observation is in line with the earlier overall glycan profile of pseudoviral gp160 of JR-CSF and BG505 SOSIP.664 trimers [43–45]. In addition, these observations are consistent with recent comparison of full-length virally-derived Env with corresponding soluble SOSIP [26,27]. In these studies, glycosylation sites displaying a mixed population of oligomannose and complex glycans in the SOSIP format were more uniformly of the complex type.

**Figure 6.**
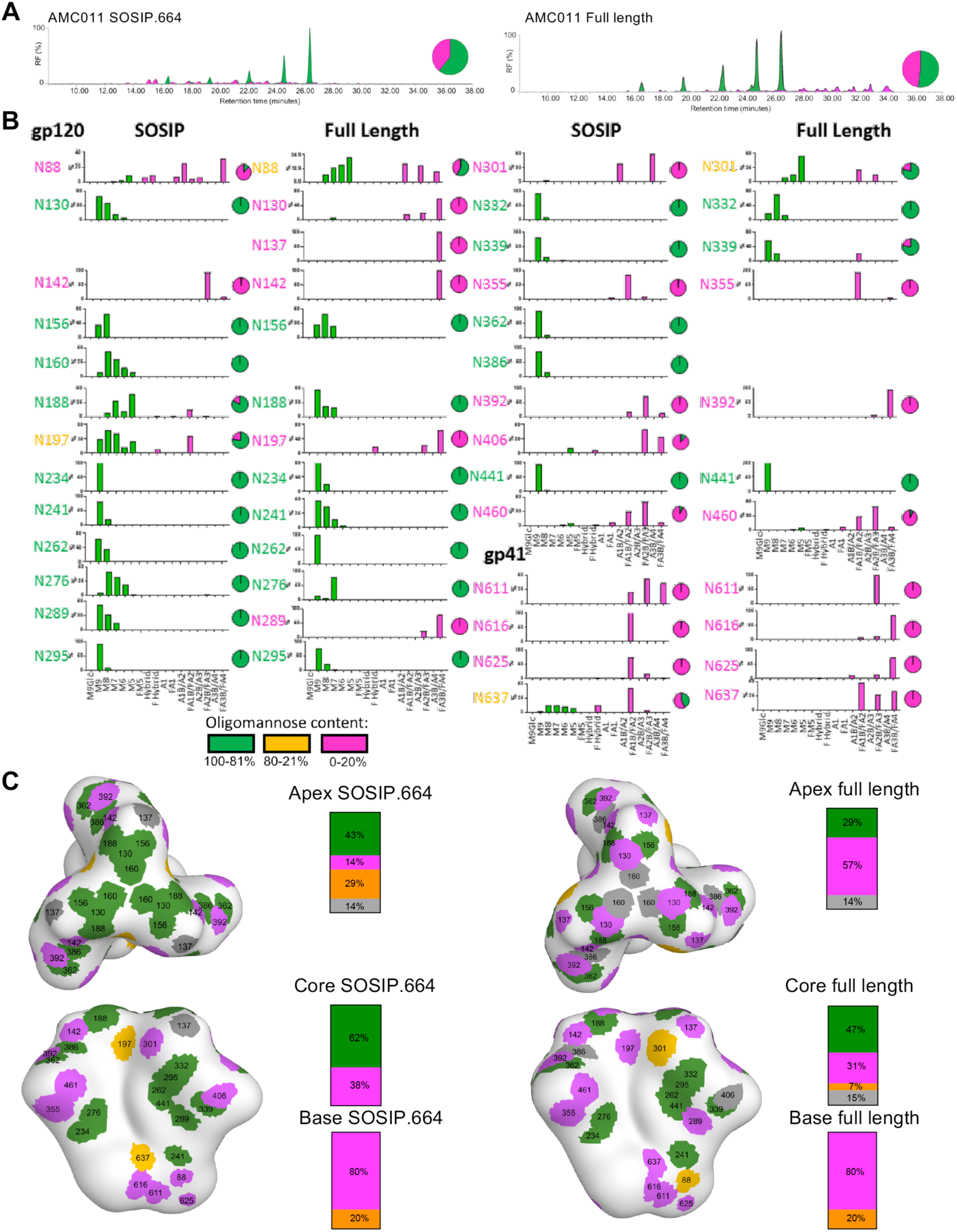
Glycan composition of full-length and SOSIP.664 AMC011 trimer. (**A**) Comparison of *N*-linked glycan profiles of full-length AMC011 trimer and SOSIP.664 Env trimer. Full-length AMC011 Env was purified in complex with PGT151 Fab. AMC011 SOSIP.664 was purified by PGT151 affinity chromatography. Green indicates oligomannose-type and hybrid glycans. Complex glycans are shown in magenta. (**B**) Relative quantification of distinct glycan types of full-length AMC011 and AMC011 SOSIP.664. The bar graphs show the relative percentage of each glycoforms at a particular site and the pie charts summarize the amount of oligomannose-type (green) and complex/hybrid (magenta) glycans on each glycan site.. (**C**) Glycan sites are colored according to their processing state in the AMC011 SOSIP.v4.2 cryo-EM map low-pass filtered to 30 Å: in green, oligomannose-type; in magenta, processed glycans; in orange, mixed glycans; in gray, glycans that could not be assigned.

To investigate the different glycan structures in more detail, we identified the *N*-glycans present in gp120 and gp41 by electrospray ionization time-of-flight mass spectrometry (ESI TOF MS) [45]. Consistent with the HILIC-UPLC data, an increase in branching and terminal elaboration of complex glycans was observed on both gp120 and gp41 in full-length compared to SOSIP.664 Env trimers (S8 Fig). Analysis of individual processed glycans showed that the full-length trimer contained a larger proportion of tri-and tetra-antennary complex glycans than the SOSIP.664 trimers (S8 Fig). The differences in mass spectrometry (MS) data between the full-length and SOSIP.664 trimers probably arise from the differences in stability and in access to glycan processing enzymes in the endoplasmic reticulum (ER). These differences might arise from at least three sources. First, SOSIP stabilization and lower propensity to sample more open conformation, might limit access of processing enzymes at specific sites. Second, glycan processing enzymes are all membrane tethered so the full-length trimer is more membrane proximal and hence more accessible to these enzymes than the soluble SOSIP.664 trimers. Third, soluble and membrane tethered Env trimers might have transit through the ER and Golgi compartments with different speeds, allowing more or less time for processing enzymes to act.

To probe the glycan structures at a site-specific level we used in-line liquid chromatography-ESI (LC-ESI) MS as previously described [45]. We were able to characterize 27 glycan sites for the SOSIP.664 trimer and 24 for the full-length trimer (Fig 6B). However, the other sites (one for SOSIP.664 and four for full-length trimer) could not be determined and/or quantified (Figs 6B and 6C). When we plotted the type of glycan at each glycosylation site onto a low-resolution map of the AMC011 SOSIP.v4.2 trimer [24], glycans at the apex and base of the Env trimer in particular, were more processed on full-length than on SOSIP.664 trimers (Fig 6C). This observation was similar to the recently described PC64 Env and the Env present on the virus [26,27,30]. However, glycans located on the core of the trimer were predominantly oligomannose-type on both trimers (Fig 6C). This echoes the differences in glycosylation observed between BG505 SOSIP.664 and virally derived BG505 gp120 [27]. We could corroborate, as previously shown by Behrens et al., the high number of oligomannose glycans across the gp120 trimer surface and the high amount of complex glycans across the gp41 surface [45,46]. However, at the individual glycan level, a number of glycans have different compositions on full-length and SOSIP.664 trimers as discussed below.

The N88 site on gp120 at the interface with gp41, contains an increased population of oligomannose glycans on the full-length compared to the SOSIP.664 trimer (Fig 6B). This may be the result of reduced accessibility to mannosidases to this membrane proximal glycan on full-length Env. At the trimer apex, the N130 site is populated by predominantly tetra-antennary complex-type glycans on the full-length trimer, whereas a large proportion of Man_9_GlcNAc_2_ moieties are present at the corresponding site on the SOSIP.664 trimer, implying that this region is somewhat protected from mannosidase digestion on SOSIP.664 but not on full-length (Fig 6B). This observation might relate the more plastic nature of the trimer apex in the full-length Env. This again recapitulates comparative glycan analyses from both BG505 and JRFL where the equivalent sites N133 for BG505 and N135 for JRFL present elevated oligomannose-type glycans in the SOSIP format [26,27]. Furthermore, near the trimer apex, the N142 site is occupied by mostly tetra-antennary complex glycans on the full-length trimer while this site contains tri-antennary glycans in the SOSIP.664 trimers. On the other hand, the N156 site is occupied by predominantly oligomannose glycans on both SOSIP.664 and full-length trimers, while the N160 site, oligomannose on the SOSIP.664 trimer, could not be assigned on the full-length trimer (Fig 6B). The same observation as for the N130 site was also made for the N289 site, where the full-length trimer is populated by tetra-antennary glycans and the SOSIP.664 trimer with oligomannose glycans (Fig 6B). A similar, but less pronounced case can be observed for N197, a glycan situated near the triad of glycans at N392, N362, and N386. This site is mixed for SOSIP.664 and processed on full-length Env (Fig 6B). The N197 site displays the same elevation in processing when BG505 is expressed in a viral context or recombinantly with a membrane tether [26,27].

The glycan sites near the CD4bs, N234, N276 and N241, are well-conserved oligomannose-type glycans in both constructs (Fig 6B). The oligomannose patch, formed by residues N295, N262, N441 and N332, is a dense region with oligomannose glycans that interact with each other and is also conserved between full-length and SOSIP.664 trimers, as might be expected (Fig 6B) [46]. Previous comparison of site-specific glycosylation of recombinant BG505 gp120 and BG505 SOSIP.664 revealed a trimer-associated mannose patch (TAMP) with up to seven sites showing elevated oligomannose levels in the SOSIP format [46]. Here, we find that some of these sites are more processed in the full-length format (e.g. N197) whereas the limited mannosidase trimming at other TAMP sites are conserved (e.g. N156 and N276). The glycan sites in gp41 showed an increased processing at all sites for the full-length trimer compared to the SOSIP.664 trimers. N637 contains some oligomannose glycans on the SOSIP.664 Env but not on full-length Env (Fig 6B). This observation is in line with the ESI MS data (S8 Fig), where oligomannose-type glycans were observed in the spectra for SOSIP.664 gp41 but not for the gp41 present on the virus [44].

Overall, full-length Env trimers contain fewer oligomannose-type glycans more complex-type glycans that are predominantly highly processed, tri-and tetra-antennary sialylated structures, compared to SOSIP.664 trimers (Figs 6 and S8).

## Discussion

We assessed the structural, biophysical and antigenic differences between the full-length AMC011 trimer and its SOSIP.664 counterpart. We purified full-length Env protein in complex with PGT145 and PGT151 Fab and we concluded that both structures showed a highly similar topology to that of published membrane-derived and SOSIP.664 trimers from various isolates, irrespective of the presence of a quaternary-specific conformational antibody. We thus show here that PGT151 binding does not induce or select for a specific Env trimer conformation that resembles the multitude of SOSIP structures, as we now show that PGT145 bound Env trimers exhibits a highly similar conformation. However, several subtle but interesting differences were found that related to glycan processing and antigenicity.

The Env protein is highly glycosylated and steric restriction of glycan processing leads to high amounts of oligomannose glycans (predominantly Man_8_GlcNAc_2_ and Man_9_GlcNAc_2_) on the outer domain of gp120 [47,48]. The data presented here compares site-specific glycan composition of full-length HIV-1 Env trimers and HIV-1 soluble immunogens and this knowledge can therefore be used for immunogen preparation since the glycans have a key role and are a component of bNAb epitopes [48]. Our glycan site-specific analysis showed that the full-length Env trimers contain higher amounts of complex glycans and a larger proportion of tri-and tetra-antennary complex types compared to SOSIP.664 trimers. The more extensively processed glycans on full-length trimers might be caused by a prolonged transit time in the Golgi apparatus compared to its SOSIP.664 counterpart, which is not membrane bound and contains smaller complex glycans, suggesting that the soluble version transits faster through the Golgi. Moreover, the differences in glycan processing may arise from the differences in accessibility to glycan processing enzymes in the ER and the Golgi, where most of the glycan processing enzymes are membrane tethered so the full-length and the soluble trimer will have different proximity and accessibility to these enzymes. Finally, the differences might arise from conformational rigidity, as the higher stability and lower propensity to sample alternative conformations of SOSIP.664 trimers might reduce accessibility of glycans to processing enzymes.

A previous report showed a comparison of soluble and full-length and ΔCT Env structures [23,30], but differences in antigenicity were not assessed due to limited protein yields. Here, we identified a high expressing full-length Env clone (AMC011) that enabled such comparisons. Based on our analyses, several differences in antigenicity could be explained by site-specific glycosylation alterations. The apparent increased conformational flexibility of the full-length AMC011 trimer, resulting in the exposure of epitopes targeted by non-NAbs, and the presence of the MPER, targeted by 4E10 and other MPER-bNAbs also demarcate key differences. Keeping the MPER domain would maximize the presentation of potent bNAb epitopes in a potential HIV-1 soluble vaccine, but whether that should come at the expense of exposing multiple non-NAb epitope remains a question.

A stable full-length Env that presents all bNAb epitopes and none of the non-NAbs epitopes could be a suitable vaccine candidate that mimics the native spike on the virus. Furthermore, such a construct may be suitable for nucleic acid delivery vectors. SOSIP.664 trimers embody many of the properties of native Env and represent the best in class trimeric Env vaccine platform currently under investigation. Here we show that subtle differences in the glycosylation may have important antigenic consequences. Moreover, the addition of the MPER epitope, as present in the full-length clone described here, may be desirable because MPER directed antibodies can be very broad and potent [49–51]. While we have now produced full-length Env in sufficient quantities for animal immunizations it may be prudent to further engineer the trimer to have a more biased antigenic profile similar to corresponding SOSIP immunogens, for example by building the SOSIP design into full-length Env trimers.

## Materials and Methods

### Design of AMC011 full-length and SOSIP.664 trimers

AMC011 is a consensus sequence from three different clonal variants of *env* genes found 8 months post-seroconversion in an elite neutralizer patient from the Amsterdam Cohort Studies on HIV-AIDS [24]. The consensus sequence was used to generate full-length AMC011 gp160 and SOSIP.664 constructs. The full-length AMC011 gp160 expression vector was created by cloning the complete sequence of *env* into the pcDNA3.1 mammalian protein expression vector. AMC011 SOSIP.664 was generated following previously published methods [4]. In short, the sequence was codon optimized for better production in mammalian cells; the A501C and T605C substitutions were introduced to form a disulfide bond between gp120 and gp41 subunits of the trimer; the I559P mutation was included to stabilize gp41; the RRRRRR cleavage site was replaced the original furin cleavage site (REKR) to enhance cleavage; the tissue plasminogen activator (tPA) signal peptide replaced the natural one to increase secretion and a stop codon at position 664 was introduced for production of the soluble protein.

### Protein production

In this study we used AMC011 SOSIP.664 and full-length protein. AMC011 SOSIP.664 was produced as previously described for other recombinant SOSIP.664 proteins [18]. In brief, HEK293F cells were transfected with a 1:3 ratio furin:Env DNA using PEImax (1 mg/mL), at a cell density of ∼1×10^6^cells/mL. Culture supernatants were harvested 7 d after transfection. SOSIP.664 trimers were purified using PGT145 affinity chromatography column as previously described [8,10]. The eluted Env proteins were further purified by size exclusion chromatography (SEC) using TBS buffer. The fractions corresponding to the trimer peak were pooled and concentrated using a MWCO concentrator (Millipore) with a 100 kDa cutoff.

PGT151 TEV IgG and PGT145 TEV IgG, in which a TEV protease cleavage site is inserted between the Fc and the Fab regions was generated and expressed in HEK293F cells as described [34,52]. In short, IgGs were expressed in HEK293F cells at 1×10^6^ cells/mL using PEImax with a ratio of 2:1 heavy:light chain. 5 d after transfection, the supernatant was harvested and passed through a 5 mL MAb select column (GE Healthcare). IgGs were eluted with 0.1 M Glycine and buffer exchanged to TBS pH 7.4.

Full-length AMC011 Env was expressed and purified as a previously published [23,30]. In brief, HEK293F cells were transfected with plasmids encoding furin and Env (furin:Env ratio of 1:3) using PEImax, at a cell density of 1.6×10^6^ cells/mL. Cells were harvested 60-65 h post-transfection. Cells were incubated with PGT151 TEV IgG or PGT145 TEV IgG and solubilized. Cell debris was centrifuged and the supernatant incubated with Protein A resin (Genscript). The protein was incubated with 0.25 mg of TEV protease/L of initial HEK293F culture and eluted with buffer containing 0.1% w/v DDM, 50 mM Tris pH 7.4, 150 mM NaCl and 0.03 mg/mL sodium deoxycholate. The same buffer was used to further purify the trimers using a Superose 6 (GE Healthcare) column. The size exclusion was run at 0.4 mL/min and fractions corresponding to the full-length AMC011 Env - PGT151 Fab complex or full-length AMC011 Env - PGT145 Fab complex were concentrated, with a MWCO concentrator to 8-9 mg/mL and 3-4 mg/mL, respectively. Protein was measured by absorbance at 280 nm using theoretical extinction coefficients calculated with Expasy (ProtParam Tool).

### Cryo-EM sample preparation

A lipid mix containing 1:1 ratio of 1,2-dioleoyl-*sn*-glycero-3-phosphocoline (DOPC) and cholesteryl hemisuccinate (CHS) was added to the concentrated Env-PGT151 Fab and Env-PGT145 Fab complex such that the final concentration was 0.14 mM. The resulting lipid-Env-Fab mixture (14 μL in total) was incubated on ice for 3 h with 15 SM-2 Biobeads (Biorad) to partially remove the deoxycholate and the DDM. 2/2 C-flat holey carbon grids with mesh size 400 (Protochips) were used for the PGT151 complex and 1.2/1.3 gold grids for the PGT145 complex (Quantifoil). All grids were plasma cleaned for 5 s using solarus advanced plasma cleaning system (Gatan) and used for the Env-PGT151 Fab and Env-PGT145 Fab sample. Blotting was performed at 4°C and 100% relative humidity using vitrobot (Thermo Fisher Scientific) as follows. First, 1 μL of 0.04% amphipol A8-35 (Anatrace) was applied to the grid followed by 3 μL of the Env sample. The grid was blotted for 3 s or 6 s for gold or C-flat grids respectively and plunged into liquid ethane. Grids were stored in liquid nitrogen until use.

### EM data collection and processing

The AMC011 full-length trimers were imaged using either the Titan Krios (Thermo Fisher Scientific), operating at 300 keV or Talos Arctica operating at 200 keV, both equipped with a K2 Summit electron detector (Gatan). The data were collected using Leginon image acquisition software [53]. Data for AMC011FL-PGT151 complex with Titan Krios was collected at 29, 000 X magnification at pixel size of 1.03 Å in the specimen plane and a dose rate of 6.763 e^-^/pix/sec for total of 8s with 250 ms frames, resulting in total dose of 51 e^-^/Å^2^. Data collected with Talos Arctica was acquired at 36, 000 X magnification at pixel size of 1.15Å and dose rate of 10e^-^/pix/sec for total of 9s with 250 ms frames resulting total dose of 68 e^-^/Å^2^.

All frames were aligned and dose weighted with MotionCor2 [54]. CTF models were calculated either with GCTF or CTFFIND3 [55]. For the PGT151 complex dataset collected with Titan Krios, 481,941 particles were picked from a total of 3,076 images using DogPicker [56], curated with 2D classification using cryoSPARC and with 3D classification using Relion 3.0 [57,58]. An ab-initio model generated with cryoSPARC was used as an initial model for 3D classification and 74,889 particles were selected for the final 3D reconstruction which was refined with soft mask to 4.2Å resolution according to the FSC 0.143 gold-standard criterion. For the data collected with Talos Arctica, the particles were sorted by 2D classification using cryoSPARC. The ab-initio reconstruction feature in cryoSPARC was used to generate an initial model and sort particles in 3D. A clean 3D class containing 51,588 particles was refined to 5.7 Å using the homogenous refinement feature in cryoSPARC.

### Determination of the approach angles of the Fabs to gp120 on the Env

To measure the differences in angles of approach of different Fabs to the gp120, we used one method that has been previously described [12]. We calculated the angle between the pseudo-2-fold axes (these axes point toward the epicenter of the epitope) and also between the axes bisecting the canonical disulfides. These two angles define the difference in the overall angular approach (rotation and tilt) of the Fab to the gp120, and then how much the Fabs rotate to interact with antigen. The sum of these two angles is roughly equivalent to the total angular difference in binding orientation calculated by the first method.

### Negative stain EM

Purified Env trimers were analyzed by negative-stain EM. A 3 µL aliquot containing ∼0.03 mg/mL of the trimer was applied for 10 s onto a carbon-coated 400 Cu mesh grid that had been glow discharged at 20 mA for 30 s, then negatively stained with uranyl formate for 30 s. Data were collected using a FEI Tecnai F20 or T12 electron microscope operating at 120 keV, with an electron dose of ∼55 e^-^/Å^2^ and a magnification of 52,000x that resulted in a pixel size of 2.05 Å at the specimen plane. Images were acquired with a Gatan US4000 CCD or Tietz TemCam-F416 CMOS camera using a nominal defocus range of 900 to 1300 nm.

### Analysis of total glycan profiles by HILIC-UPLC

N-linked glycans were enzymatically released from envelope glycoproteins via in-gel digestion with Peptide-N-Glycosidase F (PNGase F), subsequently fluorescently labeled with 2-aminobenzoic acid (2-AA) and analyzed by HILIC-UPLC, as previously described [43,45]. Digestion of released glycans with Endo H enabled the quantitation of oligomannose-type glycans.

### Assigning glycan compositions using tandem ion mobility ESI MS

The compositions of the glycans were determined by analyzing released glycans from trimers by PNGase F digestion using ion mobility MS [45]. Negative ion mass, collision-induced dissociation (CID) and ion mobility spectra were determined with a Waters Synapt G2Si mass spectrometer (Waters Corp.) fitted with a nano-electrospray ion source. Waters Driftscope (version 2.8) software and MassLynx™ (version 4.1) was used for data acquisition and processing. Spectra were interpreted as described previously [43,45]. The results obtained served as the basis for the creation of sample-specific glycan libraries, which were used for subsequent site-specific N-glycosylation analyses.

### Site-specific N-glycosylation analysis

Before proteolytic digestion, trimers were denatured and alkylated by incubation for 1 h at room temperature (RT) in a 50 mM Tris/HCl, pH 8.0 buffer containing 6 M urea and 5 mM dithiothreitol (DTT), followed by the addition of 20 mM iodacetamide (IAA) for a further 1h at RT in the dark, and then additional DTT (20 mM) for another 1 h, to eliminate any residual IAA. The alkylated trimers were buffer-exchanged into 50 mM Tris/HCl, pH 8.0 using Vivaspin columns and digested separately with trypsin, chymotrypsin and elastase (Mass Spectrometry Grade, Promega) at a ratio of 1:30 (w/w). Glycopeptides were selected from the protease-digested samples using the ProteoExtract Glycopeptide Enrichment Kit (Merck Millipore). Enriched glycopeptides were analysed by LC-ESI MS on an Orbitrap fusion mass spectrometer (Thermo Fisher Scientific), using higher energy collisional dissociation (HCD) fragmentation. Data analysis and glycopeptide identification were performed using Byonic™ (Version 2.7) and Byologic™ software (Version 2.3; Protein Metrics Inc.).

### Binding of antibodies to cell surface Env

Cell surface expression of AMC011 Env protein was assessed using flow cytometry as previously described, with some modifications [57]. HEK293F cells were transfected with plasmids encoding furin and Env (full-length AMC011 gp160) in a 1:3 ratio using PEImax at 1.75×10^6^ cells/mL. Cells were spun down 60-65 h post transfection and diluted with PBS to a final density of 10^5^ cells/mL. 10 μL of cells were incubated for 2 h with three fold serial dilutions of several monoclonal antibodies. Cells were washed twice with PBS before staining the cells with 1:50 dilution of Alexa 647-conjugated mouse anti-human IgG. Binding of MAbs was analyzed with flow cytometry as previously described [57]. Nonlinear regression curves were determined using Graphpad prism and IC_50_ values calculated.

### Neutralization assays

Neutralization assays were carried in TZM-bl cells, which express high levels of CD4, CCR5 and CXCR4 and contain luciferase genes under the control of the HIV-1 long terminal repeat promoter. We used the AMC011 chimeric molecular clone virus to perform neutralization assays. Production of the virus and neutralization assays was performed as previously described [4]. The virus was incubated with three fold serial dilutions of MAbs for 1 h. The mixture was added to the TZM-bl cells together with diethylaminoethyl. Cells were lysed ∼48 h later and luciferase activity was subsequently measured with a Glomax luminometer. IC_50_ values were calculated with the determined nonlinear regression curves using Graphpad prism.

### Bio-Layer Interferometry (BLI)

Binding studies were performed using Octet Red96 instrument (ForteBio). All the assays were performed at 500 rpm in BLI buffer (PBS supplemented with 0.01% (w/v) BSA, 0.002% (v/v) Tween 20 and 0.1% (w/v) DDM at 25°C. Anti-human IgG (AHI) probes were equilibrated in BLI buffer for 60 s. mAbs, which were diluted with kinetics buffer at a final concentration of 0.025 μg/mL, were loaded to the AHI probes for 300 s. To remove partially bound antibody to the probe, the sensors were equilibrated for 60 s with kinetics buffer. Binding to Env proteins, at final concentration of 250 nM, was determined for 300 s and dissociation for 300 s. Data analysis was carried out using Octet software and curve fitting using Graphpad prism.

### Differential Scanning Fluorimetry

Thermostability of AMC011 full-length and SOSIP.664 trimers was determined with a Nano-DSF (Prometheus). Proteins were diluted to a final concentration of 1 mg/mL. After loading 10 μL of the sample to the grade capillaries, intrinsic fluorescence signal and therefore thermal denaturation was assessed at a linear scan rate of 1°C/min with an excitation power of 15%. Unfolding transition points were detected from changes in the tryptophan fluorescence wavelength emission at 350 and 330 nm. The thermal onset (*T*_onset_) and thermal denaturation (*T*_m_) of the proteins were calculated with Prometheus NT software.

## Acknowledgements

We thank Hannah Turner, Travis Nieusma, Zachary Berndsen, Gabriel Ozorowski and Bill Anderson for assistance with microscope management and data collection. This work is supported by the International AIDS Vaccine Initiative Neutralizing Antibody Consortium, and the Bill and Melinda Gates Foundation through the Collaboration for AIDS Vaccine Discovery (OPP1084519 and OPP1115782 to A.B.W., M.C.), P01 AI110657 (A.B.W., R.W.S.), and the Scripps CHAVI-ID (1UM1AI100663 to A.B.W., M.C.). This project has also received funding from the European Union’s Horizon 2020 for Research & Innovation program under grant agreement No. 681137 (R.W.S., M.C.).

## Author Contributions

Conceptualization: ATdlP, KR, ABW and RWS. Methodology and Investigation: ATdlP, KR, CAC, JDA, MJvG, JLT. Writing – Original draft: ATdlP, KR, RWS and ABW. Writing – Review & Editing: ATdlP, KR, CAC, JDA, MvG, JLT, MC, RWS and ABW. Funding Acquisition: MC, RWS and ABW. Resources: MC, RWS and ABW.

## Data Availability

The cryo-EM maps and fitted coordinates for AMC011 FL Env bound to either PGT145 or PGT151 have been deposited to the RCSB database with accession numbers EMD-####/PDB #### and EMD-####/PDB #### respectively.

