## Supplemental Information for "Similarities and differences between native HIV-1 envelope glycoprotein trimers and stabilized soluble trimer mimetics"

#### Supporting information

### Table S1. Tier categorization of AMC011 molecular clone

| | AMC011 Tier 1B |
|---|---|
| **Sera pool** | **$ID_{50}$ (dilution) in TZM-bl cells[1]** |
| CHAVI-0642 pool | 345 |
| CHAVI-0293 pool | 724 |
| CHAVI-0598 pool | 375 |
| CHAVI-0537 | 406 |
| CHAVI-0461 | 266 |
| GMT | **399** |
| **Antibody** | **$IC_{50}$ (µg/ml) in TZM-bl cells** |
| VRC01 | 2.3 |
| 3BNC117 | 0.13 |
| CH31 | 0.22 |
| CH01 | >25 |
| PG9 | 0.12 |
| PG16 | 0.01 |
| 10-1074 | 0.01 |
| PGT128 | 0.02 |
| PGT121 | 0.03 |
| PGT151 | 0.03 |
| 2F5 | 4.3 |
| 4E10 | 7.3 |
| 10E8 | 0.66 |
| CH01-31 | 0.44 |
| 2219 | 18.55 |
| 3074 | >25 |
| 447-52D | 21.95 |
| 2557 | >25 |
| 838-12D | >25 |
| 3869 | >25 |
| 830A | >25 |
| 654-30D | >25 |
| 1570D | >25 |
| 17b | >25 |
| F105 | >25 |
| b12 | >25 |

**Table S1. Tier categorization of AMC011 molecular clone.**
[1]Values are the plasma dilution (or antibody concentration in µg/ml for the bnAbs) at which relative luminescence units (RLUs) were reduced 50% compared to virus control wells (no test sample). Lack of neutralization by the plasma samples at a 1:20 dilution is represented as a value of '10' for purposes of calculating the geometric mean titer (GMT).

### Figure S1

**A**

| | Yield (µg/L) |
|---|---|
| PC64M18C043 | ~75 |
| ZM197M | ~45 |
| BG505 | <40 |
| JR-FL | ~75 |
| AMC011 | ~350 |

**B**

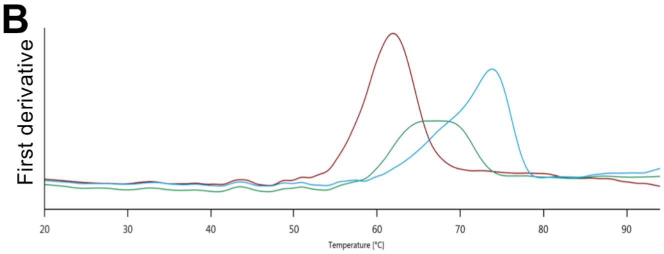


| | *Tm* onset (°C) | *Tm* Env (°C) | *Tm* Ab (°C) |
|---|---|---|---|
| SOSIP.664 | 53.5 | 61.8 | - |
| SOSIP.664 PGT145 | 58.7 | 67.0 | 70.8 |
| SOSIP.664 PGT151 | 56.5 | 73.6 | 68.2 |

**C**

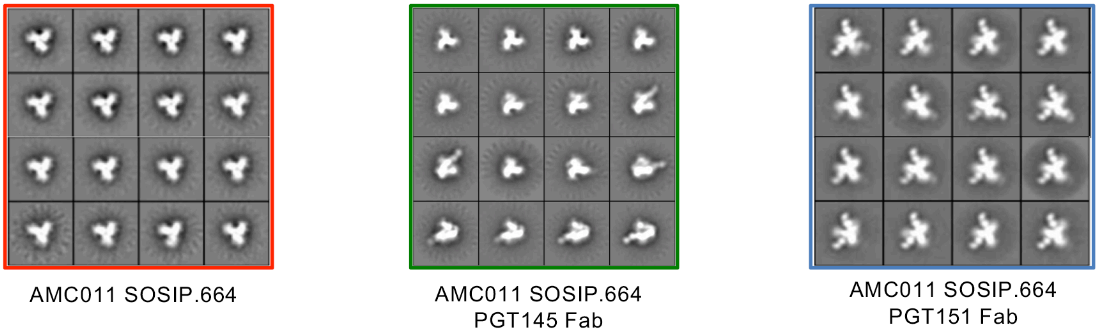


**Figure S1. Purification of full-length and SOSIP AMC011 trimer in complex with PGT151 or PGT145 Fab** (**A**) Yields obtained by purifying different full-length Env trimers. (**B**) Thermostability of the AMC011 SOSIP.664 Env trimer alone or in complex with PGT151 Fab and PGT145 Fab. The unfolding pattern of the protein complexes was determined by representing the first derivative of the curves obtained by nano-DSF. (**C**) 2D-class averages of the proteins tested in the thermostability assays in (**B**).

### Figure S2

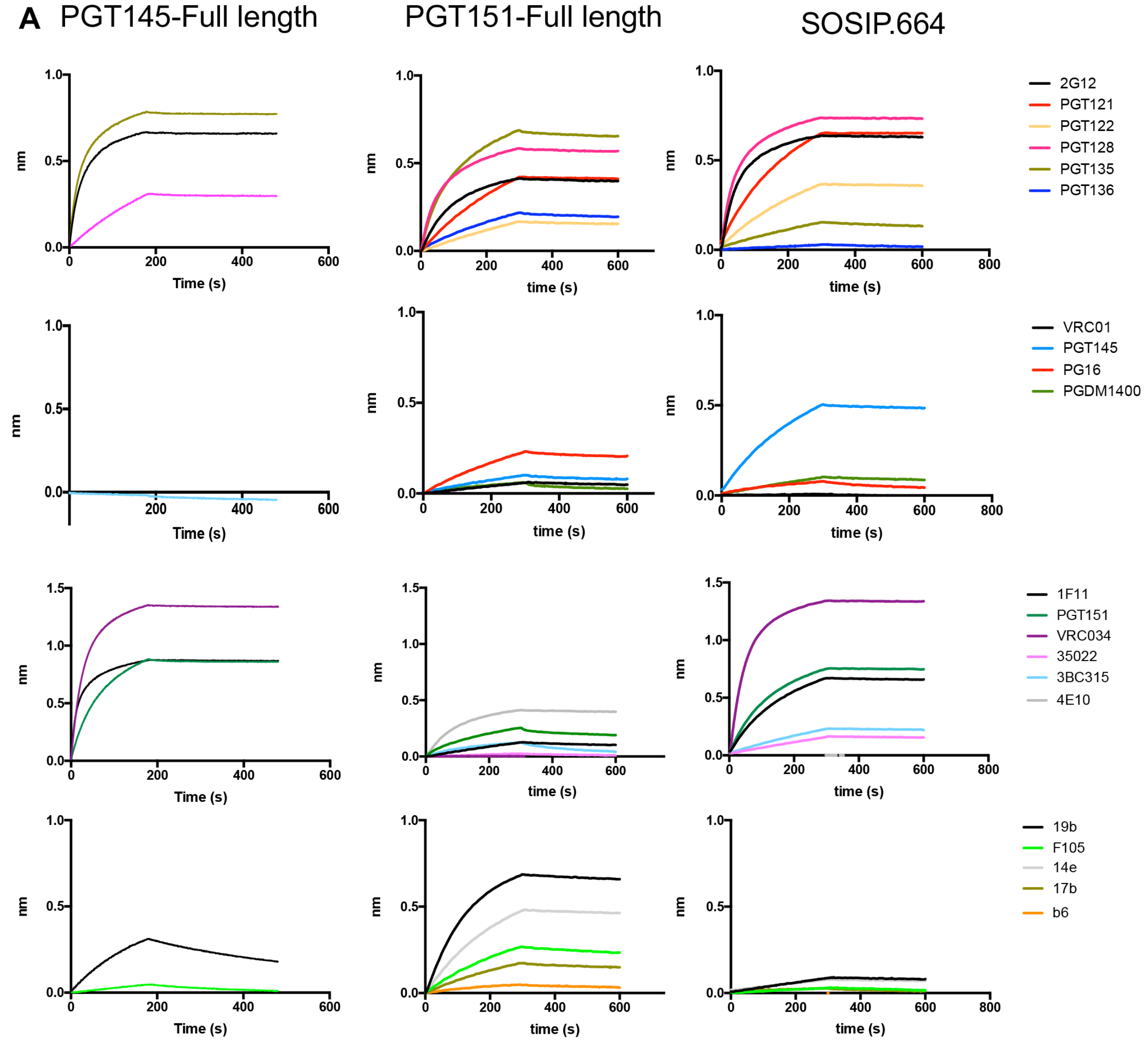


**Figure S2. Antigenicity analysis of AMC011 SOSIP.664 trimer and AMC011 full-length trimers in complex with PGT151 Fab or PGT145 Fab.** (**A**) Bio-layer interferometry (BLI, Octet) was used to determine antigenicity of AMC011 full-length and SOSIP.664 Env trimers. A panel of bNAbs and non-NAbs were immobilized to anti-human IgG sensors. AMC011 Env trimers (180 mM) were in solution through the antibody binding process. Binding curves of a panel of bNAbs and non-NAbs are shown for AMC011 full-length in complex with PGT145 Fab, AMC011 full-length in complex with PGT151 Fab and AMC011 SOSIP.664 Env trimer. The association and dissociation phases lasted between 200s and 300 s.

### Figure S3

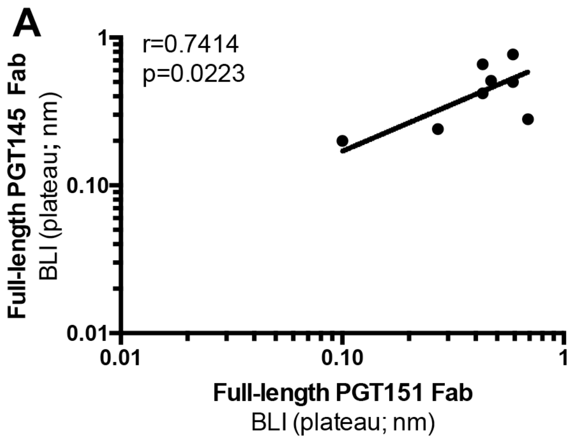


**Figure S3. Antigenicity comparison of full-length AMC011 Env trimer bound to PGT151 Fab and bound to PGT145 Fab by BLI.** Antibodies that compete with the Fabs bound to the full-length trimers, because of epitope overlapping, were omitted (PG16, PGDM1400 and PGT145 for Env bound to PGT145 Fab, and VRC34, ACS202, 3BC315 and 35O22 for Env bound to PGT151 Fab). The Spearman correlation coefficient is shown.

### Figure S4

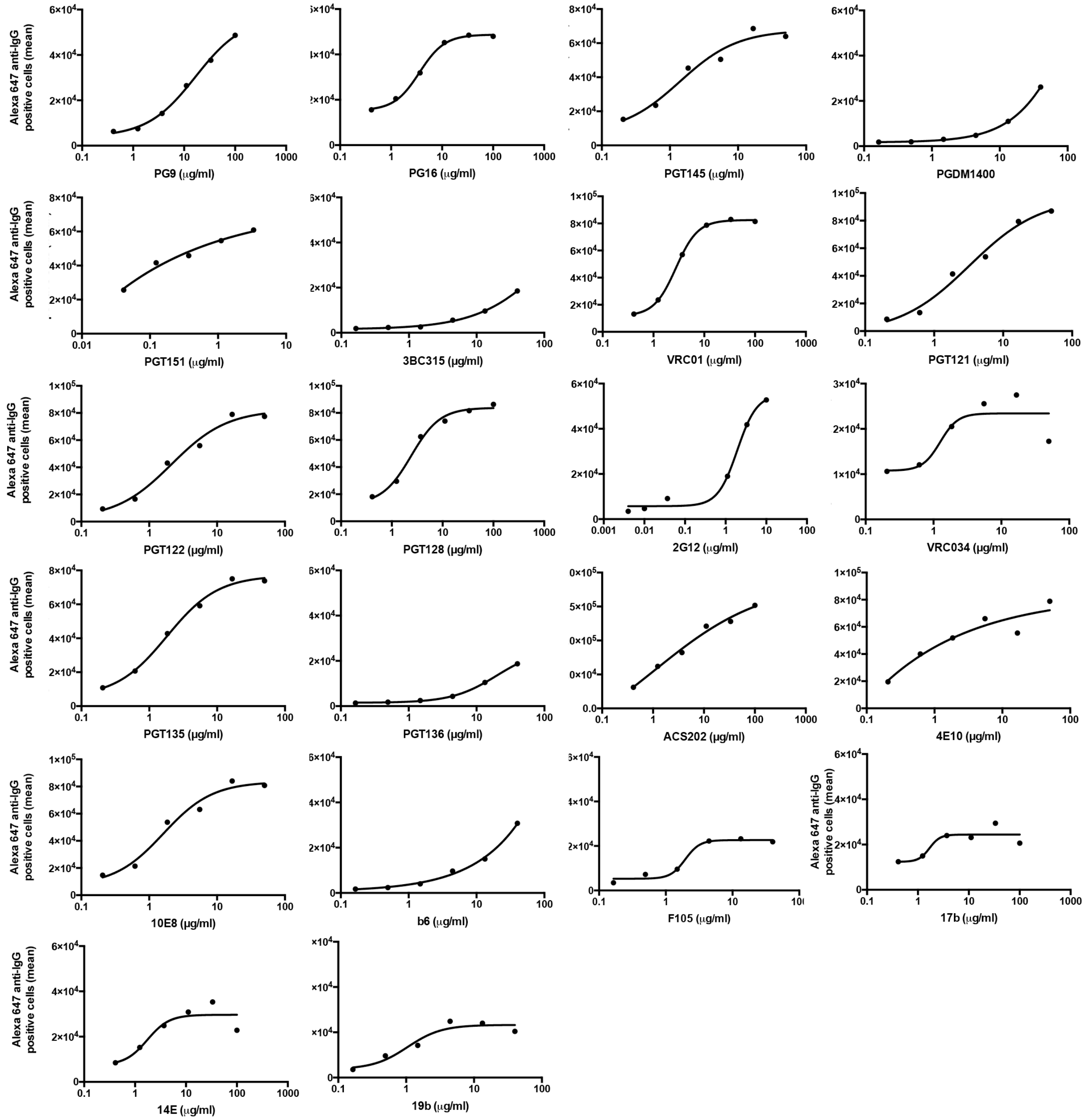


**Figure S4. Binding of a panel of bNAbs and non-NAbs to AMC011 trimers expressed on the surface.** Flow cytometry was used to determine binding of escalating concentrations of bNAbs and non-NAbs to AMC011 expressed on the surface of HEK 293F cells. Data are a representative for four different experiments.

### Figure S5

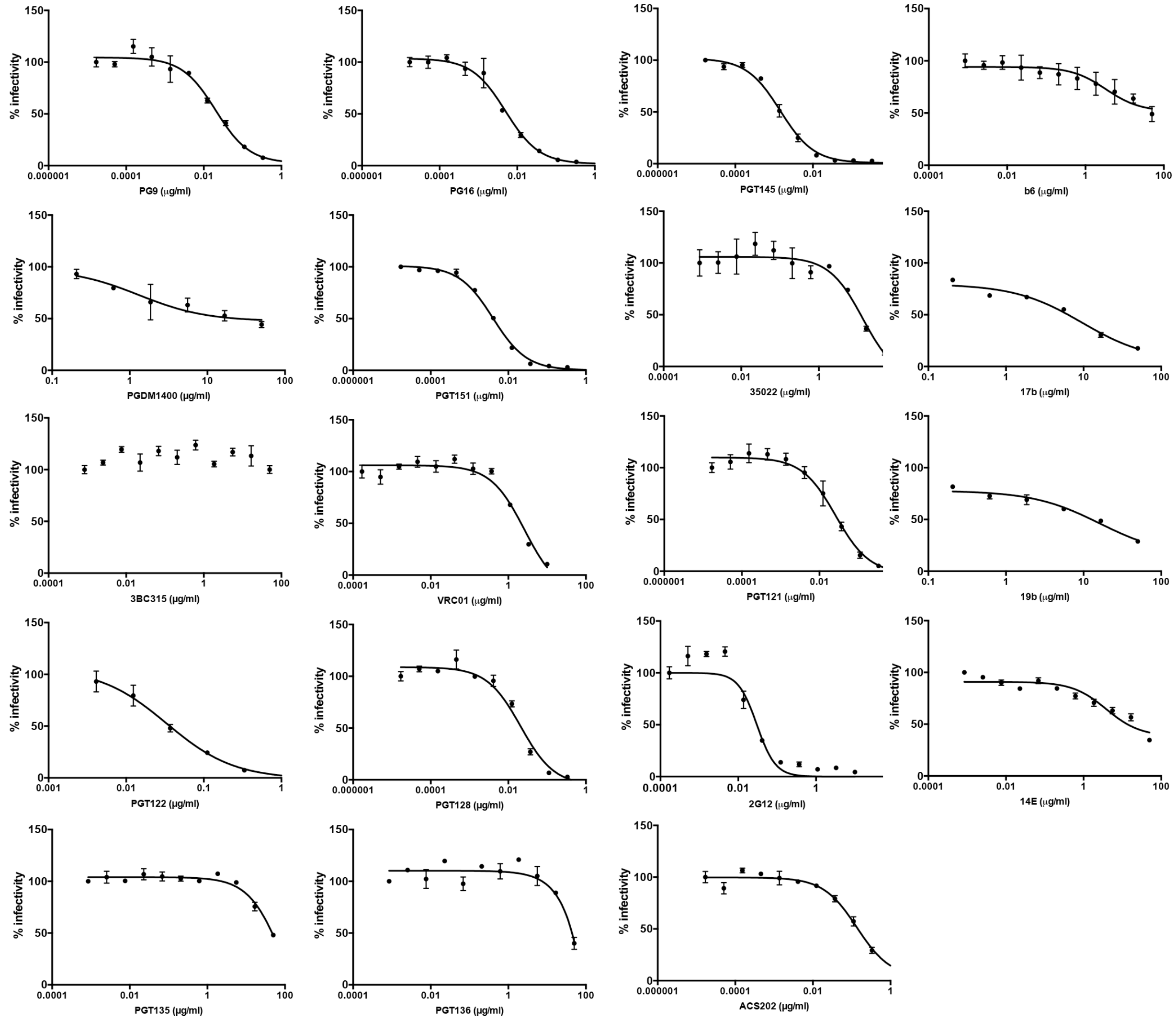


**Figure S5. Neutralization of the AMC011 molecular clone by a panel of bNAbs and non-NAbs.** TZM-bl neutralization assays were performed a minimum of two times generating duplicate titration curves in each experiment.

#### Figure S6

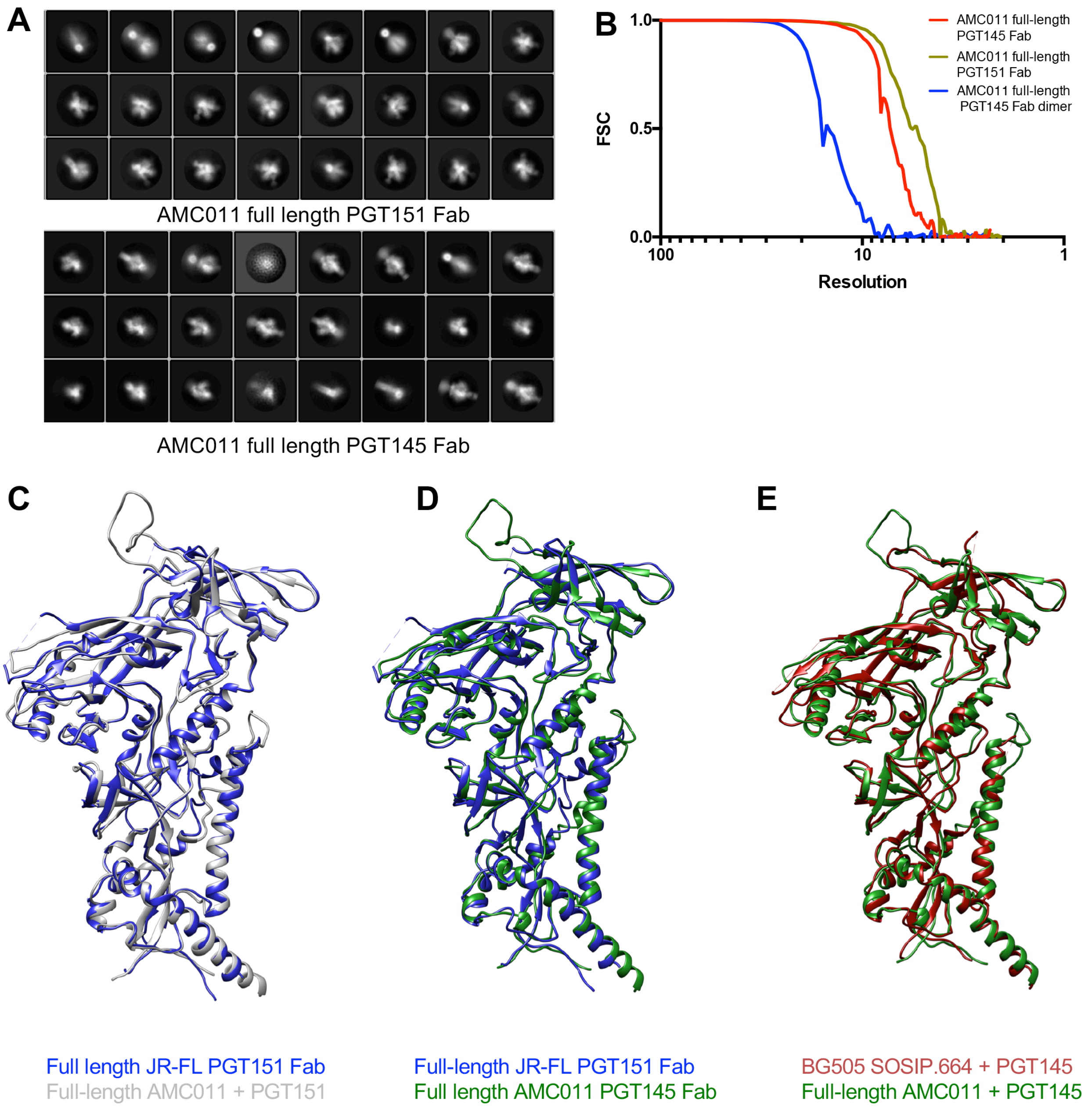


**Figure S6. Single particle analysis of the full-length AMC011 Env trimer in complex with PGT151 Fab or PGT145 Fab.** (**A**) 2D class averages of the full-length AMC011 Env in complex with PGT151 Fab (upper panel) and with PGT145 Fab (lower panel). (**B**) Fourier shell correlation (FSC) curves of the reconstructions of the full-length AMC011 Env in complex with PGT151 (green) and in complex with PGT145 Fab (trimer in red; dimer in blue). Superimposition of the JR-FL ΔCT trimer bound to PGT151 Fab (in blue) with: (**C**) full-length AMC011 bound to PGT151 Fab (in gray) or (**D**) PGT145 Fab (in green). (**E**) Superimposition of the full-length AMC011 Env bound to PGT145 (colored in green) and BG505 SOSIP.664 bound to PGT145 (colored in red; PDB:5V8L).

#### Figure S7

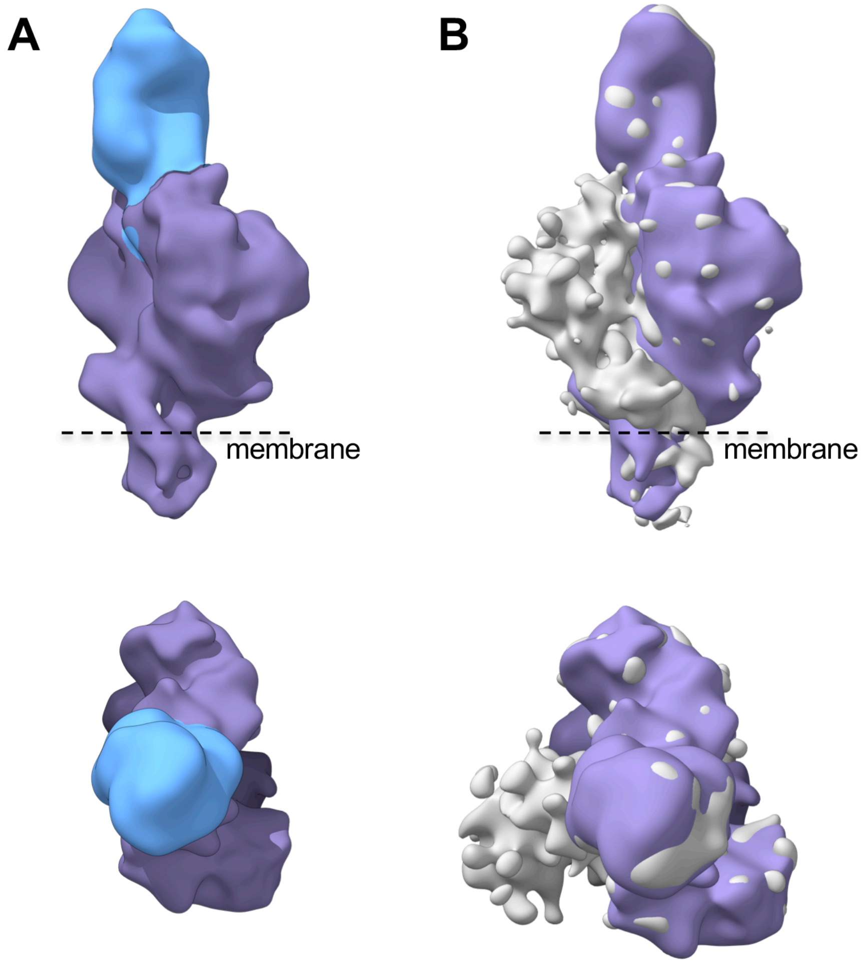


**Figure *S*7**. **Cryo-EM structure of full-length AMC011 Env bound to PGT145 Fab dimer.** (**A**) Side and top views of the cryo-EM reconstructions of full-length AMC011 Env dimer bound to PGT145 Fab. (**B**) Superimposition of the cryo-EM maps of full-length AMC011 trimer and full-length AMC011 dimer bound to PGT145 Fab.

**Figure S8**

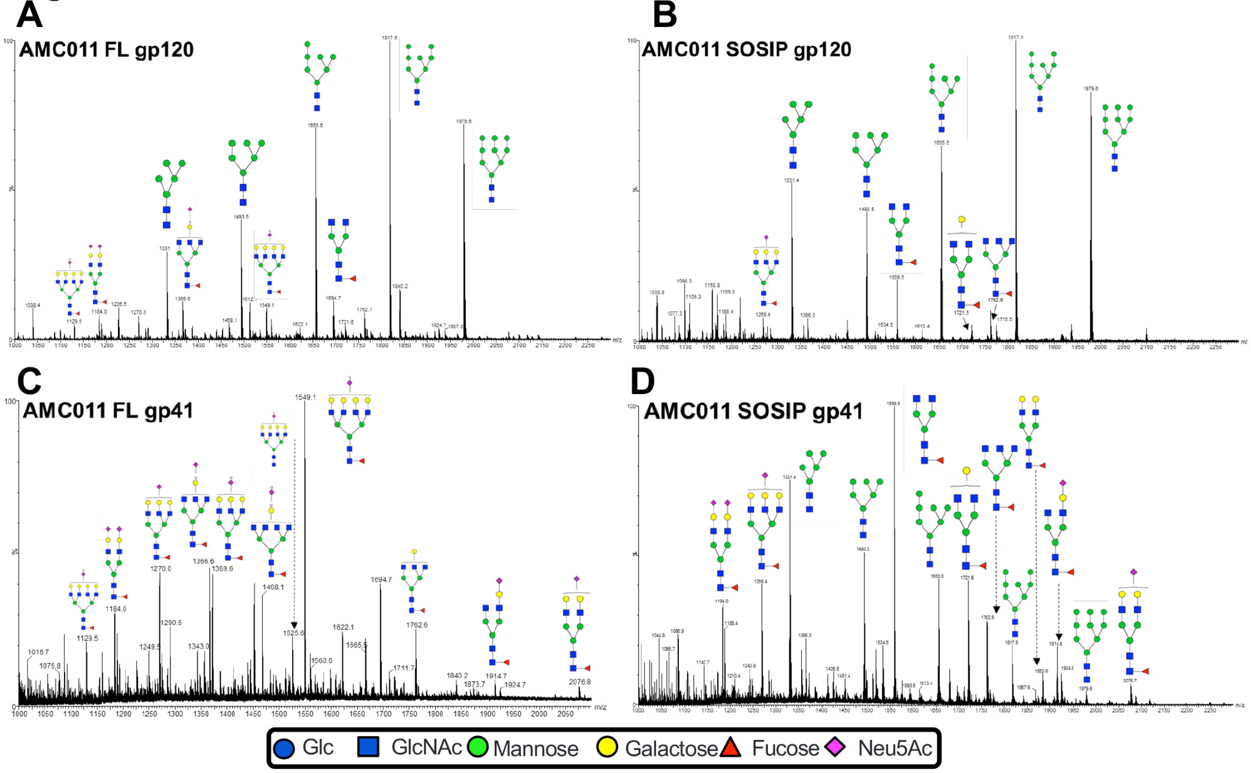


**Figure S8. Negative ion ESI-IM TOF MS analysis of glycans released by PNGaseF.** Negative-ion electrospray spectrum of N-linked glycans extracted from AMC011 full-length gp120 (**A**), AMC011 SOSIP gp120 (**B**), AMC011 full-length gp41 (**C**) and AMC011 SOSIP gp41 (**D**). Prominent peaks are annotated with the corresponding compositions, using Consortium for Functional Glycomics symbolic nomenclature.
